# A multimodal interrogation of Broca’s area in the pediatric and adult human brain

**DOI:** 10.64898/2026.07.22.739934

**Authors:** Juan Moriano, Tanzila Mukhtar, Jimmy Tsz Hang Lee, Fani Memi, Jonathan Augustin, Rachel Leonard, Joseph Asfouri, I-Ling Lu, Clara Siebert, Jennifer Ja-Yoon Choi, Guolong Zuo, Yinshui Chang, Stanislaw Makarchuk, Elina Jin, Martin Prete, Shaohui Wang, Qiuli Bi, Yuhan Hao, Liz Tuck, Katy Tudor, Vanessa B. Pereira, Hon Man Chan, Jasmine Halliwell, Benjamin Rumney, Holly Anderson, Arren Ramsey, Fabian Theis, Mercedes Paredes, Xianhua Piao, Elizabeth E. Crouch, Arturo Alvarez-Buylla, David Rowitch, Eric J. Huang, Tomasz J. Nowakowski, Omer Ali Bayraktar, Arnold Kriegstein

## Abstract

Language is a defining trait of our species, and disruptions in language acquisition can have profound consequences to the individuals affected. Uncovering the neurodevelopmental basis of this complex trait requires detailed molecular and cellular insights into the neocortical areas that support linguistic abilities. Here we performed joint gene expression and chromatin accessibility profiling at single-nucleus resolution (10x Genomics Single cell Multiome) and spatial transcriptomic profiling (Xenium high-plex *in situ* spatial transcriptomics) of Broca’s area alongside adjacent motor cortical areas. We profiled individuals from different ancestries (European and African) and developmental stages (infancy, childhood, adolescence, and adulthood). We provide a high-resolution dissection of the cellular and molecular architecture of Broca’s and motor cortical areas across early life stages and anchor the trajectories to the cellular states found in the adult human brain. We identify distinct area- and stage-specific cellular signatures, including a prominent role of glia populations and interneuron subtypes contributing to cytoarchitectonic specializations. Using longitudinal single cell spatial transcriptomic profiling, we orthogonally validate our consensus cell taxonomy and spatially resolve layer enrichment of neuronal and astrocyte subtypes that distinguish Broca’s area and motor cortex. We also uncover cell type-specific molecular signatures that distinguish cell developmental trajectories in these cortical areas, including an early molecular code established by differential expression of cadherin genes that might contribute to area-specific intercellular communication. We also identify cell type-specific vulnerabilities to language-related neurodevelopmental and neuropsychiatric disorders, with selective susceptibility of particular somatostatin-positive interneuron subtypes to ASD/ADHD. Finally, evolutionary analysis of differentially accessible regions between Broca’s area and motor cortex suggests that genetic mutations that might have contributed to the emergence of linguistic abilities accumulated over the course of million years following the divergence of human and chimpanzee lineages. Together, our study provides a comprehensive molecular, cellular and spatial definition of Broca’s area and motor cortex, laying the groundwork for investigations into unique aspects of human cognition and related neurodevelopmental and neuropsychiatric disorders.

## INTRODUCTION

The faculty of language has long been regarded as one of the most distinctive characteristics of our species. As noted by Charles Darwin, compared to other species, humans possess an “infinitely larger power of associating together the most diversified sounds and ideas”^1^. Disorders affecting language impairments can compromise educational attainment^2^ and quality of life, posing a significant lifelong burden^3^. The neuroanatomic infrastructure underlying language processing in the human brain comprises a distributed network of cortical areas from both hemispheres as well as subcortical regions^4,5^. Among these, Broca’s area occupies a position of central prominence. In 1861, Paul Broca described a patient with severe impairments in language production and identified the left inferior frontal lobe as the speech center of the human brain^6^. Subsequent landmark studies crucially informed by lesion-deficit clinical cases, including those of Wernicke^7^ and Lichtheim^8^, supported a (classical) model of language function where Broca’s area in the inferior frontal gyrus, Wernicke’s area in the superior temporal cortex, and white matter connections linking these areas, serve as core neurobiological substrates for language production and comprehension^9^. These seminal studies and ensuing efforts rendered tractable not only the investigation of neural substrates of high order cognitive abilities, but the interrogation of molecular and cellular determinants of neurological diseases.

Broca’s area, critically responsible for language production, has been classically divided into Brodmann area (BA) 44 and 45 (and to some extent BA47)^10,11^, located adjacent to primary motor (BA4) and premotor (BA6) cortex in the ventrolateral prefrontal cortex (VLPFC). In comparison to primary motor areas, large pyramidal cells in cortical layer III and a relatively developed layer IV are among the most conspicuous cytoarchitectonic features of Broca’s area^12^. In addition, an increased proportion of neuropil to cell body ratio adds to the microstructural distinctiveness of Broca’s area^13^. Although limited by regional specificity, temporal coverage, cellular resolution and/or sample size, previous studies have nonetheless uncovered transcriptomic signatures associated with Broca’s area^14–17^. Notably, layer-specific gene expression differences between the inferior frontal gyrus and superior temporal cortex have implicated *SLIT* genes, which have been linked to cortico-cortical connectivity and dyslexia^16^. These distinctions further extend to protein expression levels of neurotransmitter receptors contributing to the spatial molecular complexity of BA44 and BA45^18,19^.

Notwithstanding these observations, the molecular and cellular specializations of Broca’s area and of cortical areas involved in linguistic abilities remain poorly understood. This effort is also impeded by limited access to pediatric brain samples, which substantially hinders investigation into the ontogeny of brain networks specialized for language processing.

In this study, we interrogate molecular, cellular and cytoarchitectonic features that define Broca’s area in comparison to motor cortex across postnatal stages of brain development. We leverage the joint profiling of gene expression and chromatin accessibility paired with spatial transcriptomics to establish a comprehensive developmental atlas and create a high-resolution taxonomy of cell types across language- and motor-related areas. We uncover molecular and cellular dynamics underlying area- and stage-specific developmental trajectories, revealing significant differences in the abundance of astrocytes, oligodendrocyte progenitor cells (OPCs) and specific interneuron subtypes. We further identify an early molecular code, established by differential expression of type II cadherin genes during infancy, that is predicted to modulate intercellular communication between glutamatergic and GABAergic neurons distinguishing Broca’s area and motor cortex. We provide a comprehensive spatial transcriptomics profiling of cortical cytoarchitecture with high-resolution cell type mapping across areas and developmental stages. We find area-specific enrichment of particular glutamatergic and GABAergic neuronal subtypes that validate, and extend, our 10X multiomics analysis. We observe area-specific differences among interneurons across both upper- and deep-layer subtypes as well as astrocytes that are already evident at pediatric ages, likely reflecting early mechanisms of cortical arealization. We additionally uncover cell type-specific differential expression of candidate disease genes implicated in speech and/or motor impairments, including a calcium-dependent activator protein, *CADPS2*, that might have been under evolutionary pressures in modern humans. By integrating cell type–resolved chromatin accessibility profiles with high-risk genetic variants, we identify selective cellular correlates of neurological disorders affecting language and motor function, most notably a susceptibility of particular somatostatin-positive interneuron subtypes to ASD/ADHD during early brain development. Finally, our evolutionary analysis of regulatory regions that differ in accessibility profiles between Broca’s area and motor cortex suggests that genomic specializations contributing to the emergence of linguistic abilities accumulated over millions of years following the divergence of human and chimpanzee lineages.

## RESULTS

### A multimodal cell atlas of Brodmann’s areas involved in language and motor functions

To comprehensively characterize the diversity and molecular profiles of cell types in Broca’s area, as well as motor/premotor cortical areas, we performed 10X Genomics Multiome profiling to simultaneously capture gene expression and chromatin accessibility profiles at single cell resolution (Figure 1A). We profiled a total of 41 samples from 11 neurotypical individuals (6 male, 5 female). Broca’s area (BA) was micro-dissected to encompass Brodmann (BA) areas BA44 and BA45, and adjacent motor cortex to include primary motor (BA04) and premotor (BA06) areas (all from left hemispheres; Figure S1 and Table S1). We densely profiled early pediatric developmental stages that are most relevant to the development of linguistic abilities (spanning the transition infancy to childhood, from 0 to 6 years), as well as the adult brain (25-36 years). After stringent quality control, we retained 210,966 high-quality nuclei, with a median of 1,922 genes, 3,696 transcripts, and 7,851 ATAC fragments detected per nucleus (Figure S1C). Data from all samples were found to integrate well into a low dimensional embedding following data integration and construction of a weighted nearest neighbor graph that combined both RNA- and ATAC-sequencing data modalities (Figure 1B and S1D-F).

**Figure 1.**
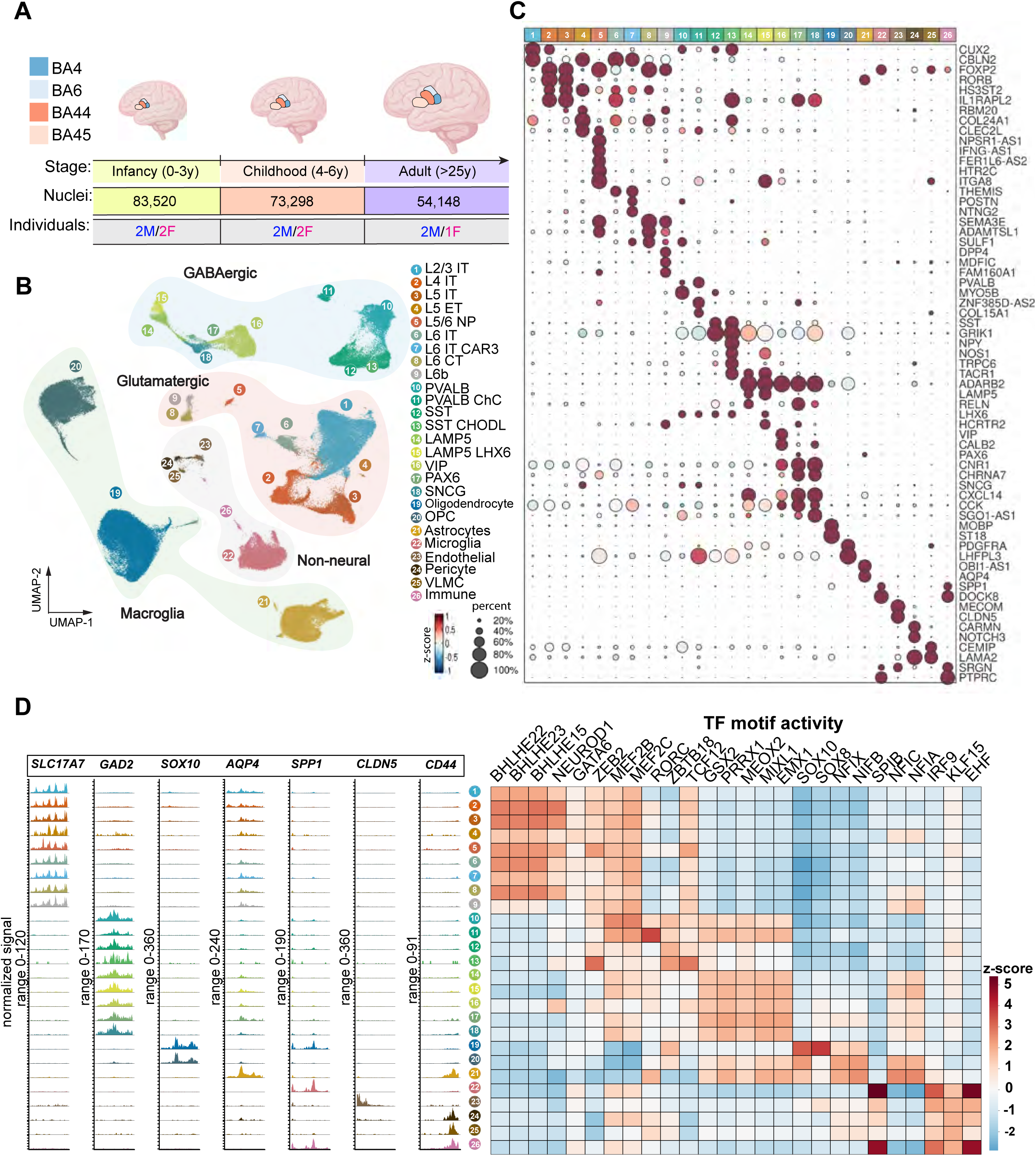
(A) Summary of experimental design of this study. (B) UMAP embedding for the snMultiome (gene expression and chromatin accessibility) data, highlighting major classes and subclasses. (C) Marker gene expression across the 26 subclasses. (D) Chromatin accessibility profiles for selected marker genes along with inferred transcription factor motif activity across same subclasses.

We establish a comprehensive multimodal atlas spanning language- and motor-related cortical areas across development and provide a high-resolution consensus cell taxonomy generated through robust, hierarchical iterative clustering with stringent criteria for marker gene differential expression and batch correction (see Methods). The multimodal dataset was first resolved into three main cell classes: Glutamatergic (n=55,117 cells), GABAergic (n=37,479 cells), and non-neuronal/non-neural cells (n=118,310 cells), with 26 subclasses (Figure 1 B-C). Subclasses include glutamatergic neurons within specific cortical layers (L2-6), GABAergic neurons defined by well-established marker gene expression (*SST, PVALB, VIP, SNCG, PAX6, LAMP5 and LAMP5 LHX6*) and non-neuronal/non-neural cells identified as macroglia cells (oligodendrocytes, OPCs and astrocytes), and other non-neural cells (microglia, vascular endothelial cells and pericytes, leptomeningeal cells, and immune cells). Gene expression and chromatin accessibility profiles as well as inferred transcription factor motif activities recapitulate established markers for these subclasses (Figure 1C-D). Oligodendrocytes were the most abundant subclass in our dataset (20.72%), followed by layer 2-3 intratelencephalic neurons (L2/3 IT; 14.33%), OPCs (13.57%) and astrocytes (10.67%) (Table S2). At a finer level of resolution, we identified 69 distinct clusters (n = 44 GABAergic, 15 Glutamatergic, 10 Nonneuronal/Non-neural) which were largely conserved across Brodmann areas and developmental stages (all donors were found well represented across clusters) (Figure 2A and S2-6; Table S3; see Methods). Consistent with previous reports^20^, GABAergic subclasses exhibited a marked molecular diversity, most prominently within the SST (n = 13) and VIP (n = 10) subclasses. We subsequently validated the laminar distribution of our taxonomy clusters with spatial transcriptomics (see below).

**Figure 2.**
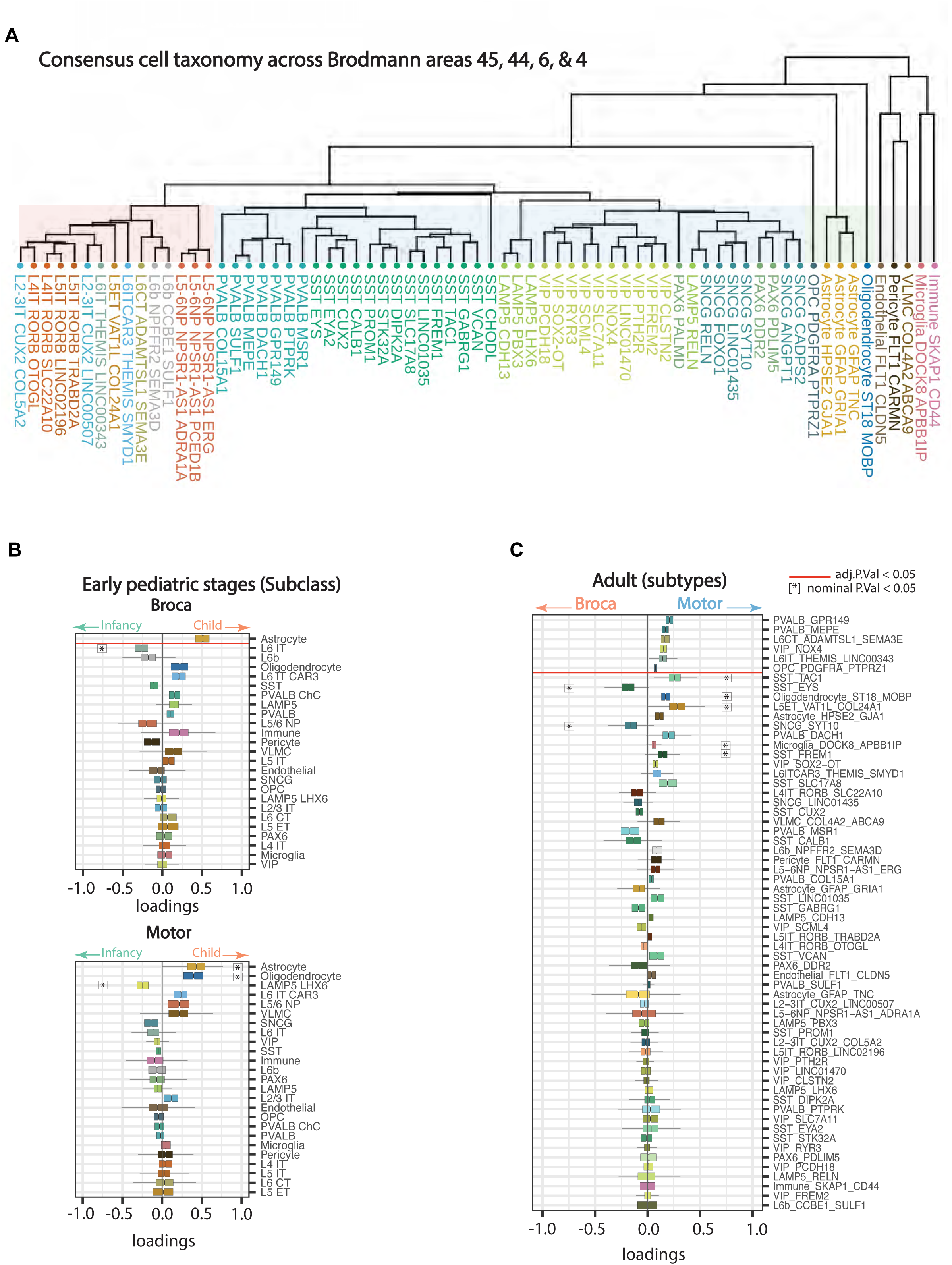
(A) High resolution consensus cell taxonomy across Brodmann areas. (B) Differential cell abundance at early pediatric stages in either Broca’s area (top) or motor cortex (bottom) at the subclass level. Statistical significance was considered if adjusted p-value after multiple comparisons was below 0.05. (C) Differential cell abundance between Broca’s area and motor cortex in adulthood, at cell taxonomy level. Significant results at adjusted p-value below 0.05. Sample sizes for Broca (BA44/45) and Motor cortex (BA06/04): Broca at adult: n = 6, Motor at adult: n = 5, Broca at childhood: n = 8, Motor at childhood: n = 7, Broca at infancy: n = 8, Motor at infancy: n = 7.

### Cell compositional changes contributing to cytoarchitectonic specializations of Broca’s area and motor cortex

Cytoarchitectonic specializations might entail developmental stage-dependent differences in cell abundance associated with the functional maturation of cortical areas. To test for significant differences in cell proportions across time as well as between Broca’s area (BA44/45) and motor cortex (BA06/04), we performed a compositional data analysis based on an isometric log-ratio transformation of cell type proportions with a resampling strategy to evaluate the significance of between-groups differences^21^ (see Methods). We detected a statistically significant increase of astrocytes in Broca’s area in the transition from infancy to childhood (adj. p-value < .05), as well as a trend toward increases of astrocytes and oligodendrocytes in motor cortex (nominal p-value < .05; Figure 2B; Table S4). Consistently across cortical areas, we observed postnatal expansion of oligodendrocytes, together with a significant increase in the OPC population in Broca’s area during childhood in comparison to adulthood (adj. p-value < .05; Figure S7A; Table S4). At the level of taxonomy clusters, dynamic changes in cell abundance were generally modest and suggested that the increase in astrocyte abundance was primarily driven by two astrocyte subtypes (*GFAP*+*GRIA1*+ and *HPSE2*+*GJA1*+; nominal p-value < 0.05; Figure S8 and Table S4). The dynamic increase in abundance of these glial populations coincides with major developmental changes in VLPFC, including the strengthening and maturation of white matter fiber tracts projecting to these areas during infancy and childhood^22,23^. These findings position the spatiotemporal control of glial population dynamics as a significant component of developmental programs in the early pediatric VLPFC.

Next, we assessed area-specific differences in cellular composition at each developmental stage. Overall, we detected only modest differences in cell type abundance between Broca’s area and adjacent motor cortical regions at the subclass level. Consistent across developmental stages and in agreement with established cytoarchitectonic distinctions, L4 IT and L5 ET glutamatergic neurons showed a trend towards increased relative abundance in Broca’s area and in motor cortex, respectively (nominal p-value < .05; Table S4; Figure S7B). By leveraging the higher granularity afforded by our consensus cell taxonomy, we observed the most pronounced changes in cell abundance in adults, while modest differences were detected at early pediatric stages (Figure 2C and S7C; Table S4). We found statistically significant increases in subtypes of glutamatergic L6 neurons (*L6 CT ADAMTSL1 SEMA3E* and *L6 IT THEMIS LINC00343*), as well as PVALB and VIP interneurons (*PVALB GPR149; PVALB MEPE; VIP NOX4*) and OPCs (adj. p-value < .05; Figure 2C and S7C) in motor cortex in comparison to Broca’s area. Only two interneuron subtypes, *SST EYS* and *SNCG SYT10*, were enriched in Broca’s area relative to motor cortex (nominal p-value < 0.05; Figure 2C; Table S4). These findings suggest that cytoarchitectonic specializations in VLPFC and motor cortex extend beyond differences in layer 4/5 glutamatergic neurons and additionally involve specific interneuron subtypes.

### Single cell resolution spatial transcriptomic maps of Broca’s area and motor cortex

To spatially resolve the molecular and cellular states we previously defined, and to compare cytoarchitectonic features of Broca’s area and adjacent motor cortex, we used Xenium spatial transcriptomics (ST) technology for probe-based imaging of single cell transcriptomic profiles *in situ*. We profiled 4 donors, spanning infancy (5 months), childhood (4 years), adolescence (9 years) and adulthood (36 years), encompassing Brodmann areas BA44, BA45, and BA06. We serially sectioned tissue blocks to generate two complementary Xenium ST datasets (Figure 3A). First, we curated a 460-plex gene panel encompassing subclass-, area- and developmental stage-specific markers as well as genes involved in cell-cell communication based on our 10X Multiome dataset (see below). This panel was the basis of our main ST dataset, which comprised over 3 million cells across 56 tissue sections following quality control. Second, we applied the 5,000-plex Xenium Prime probe panel to a single section per sample, yielding 921,487 cells across 12 sections, which were used to validate and augment our primary ST dataset. After stringent quality controls (Figure S9), we annotated our Xenium ST datasets via label transfer from our age- and region-matched 10X Multiome consensus cell taxonomy (Methods, Figure S10A-B). The marker gene expression, hierarchical clustering and cortical layer identity of these cell types was consistent between 10X Multiome and Xenium datasets (Figure 3B,C, Figure S10A-B), validating our high-resolution consensus cell taxonomy.

**Figure 3.**
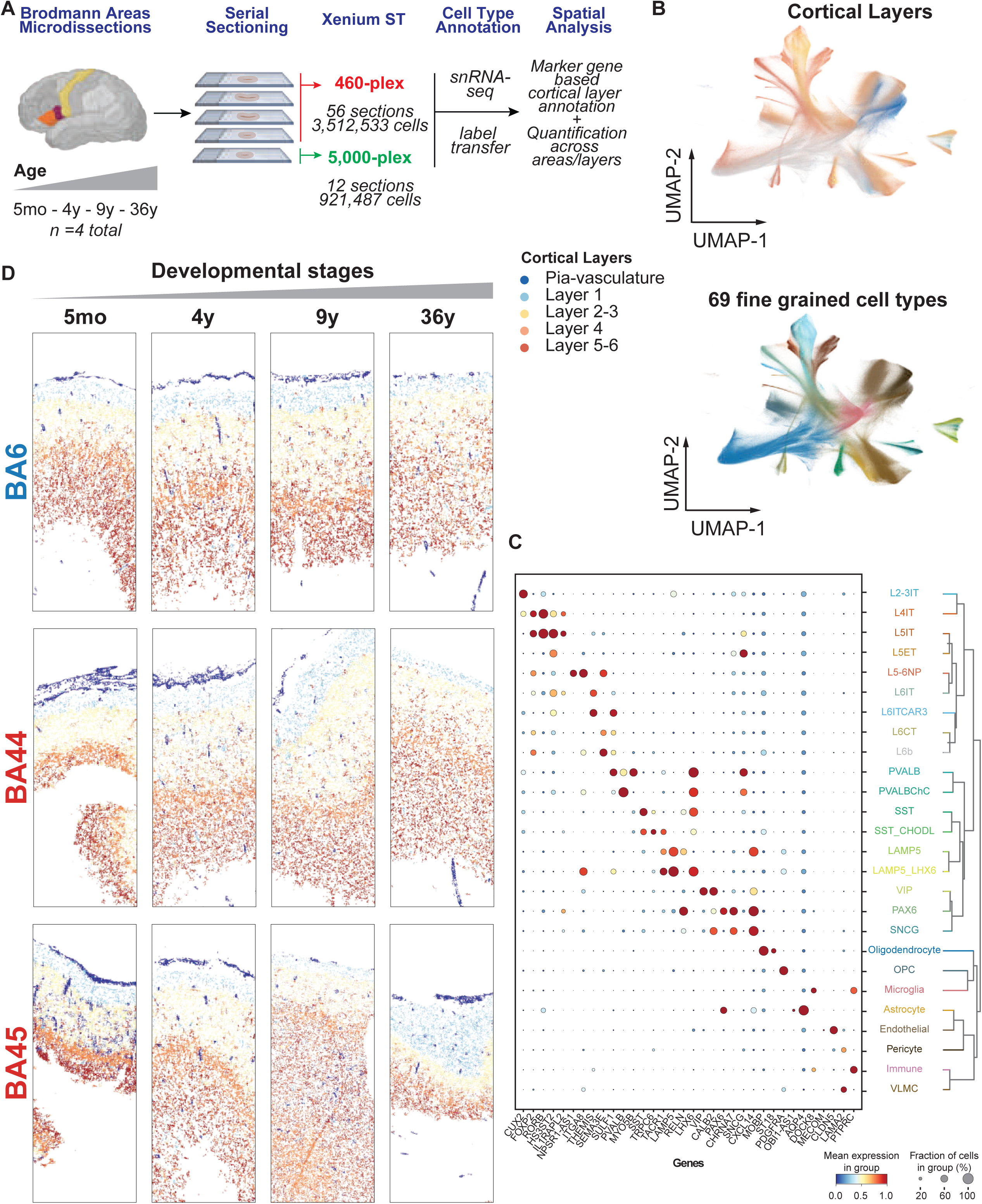
(A) Single-cell spatial transcriptomics workflow for profiling Broca’s area and premotor cortex. Below panels show Xenium 460-plex ST data unless indicated. (B) UMAP embeddings for ST data indicating our fine-grained cell type annotation (bottom) and cortical layer identities assigned based on layer-specific gene signatures (top). (**C**) Dotplot showing marker gene expression across 26 annotated cell subclasses in ST data. (D) Spatial maps of select tissue sections across Brodmann areas and developmental stages. Cells are colored according to assigned cortical layer identities.

We used three complementary approaches to resolve the cortical cytoarchitecture of Broca’s area and premotor cortex across our ST data. First, we annotated the cortical layer identities of cells based on their expression of known layer-specific gene signatures (Methods). This analysis broadly recapitulated cortical layers, delineating L1, L2-3, L4 and L5-6, as well as the pia-vasculature and white matter compartments based on their laminar position and cellular composition across cortical areas and developmental periods (Figure 3B,D). An orthogonal spatially-aware clustering approach provided consistent results^24^ (Methods, Figure S10C). Second, we quantified the distribution of cell types along cortical depth, which we normalized into bins between the pial surface and white matter (Figure 4A). Finally, we quantified differential cell type abundance between Brodmann areas per each donor / developmental period (i.e. different tissue sections per block treated as replicates) (Figure 4B). Given that our experimental design was limited to one individual per developmental stage, we defined area-enriched cell types as those with significant differences across multiple developmental stages.

**Figure 4.**
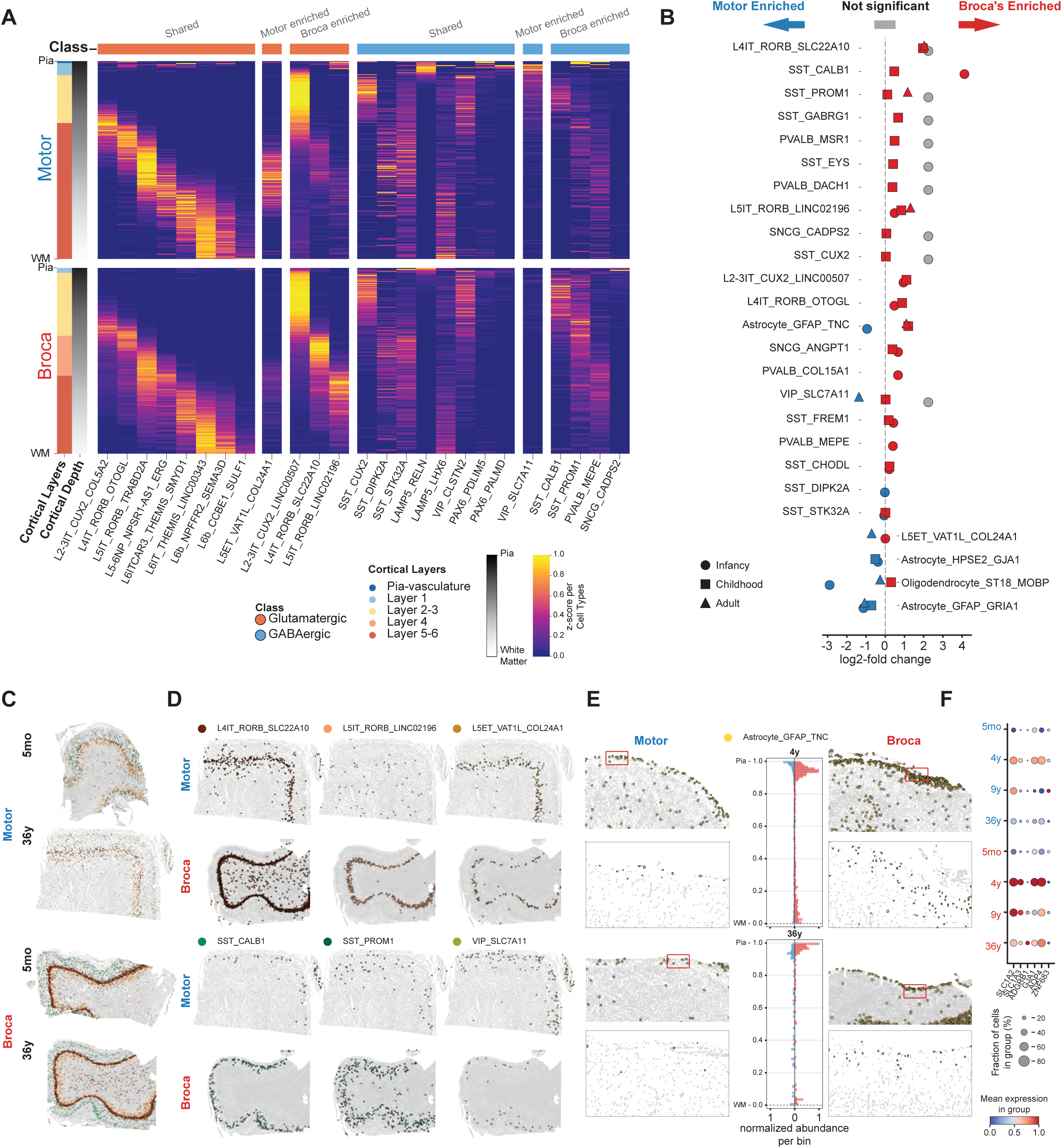
(A) Heatmaps showing normalized cell type abundances across premotor (BA06) and Broca’s area (BA45) that are binned across cortical depth (200 bins) from pia-vasculature (top) towards white-gray matter boundary (bottom). (B) Differential cell abundances between BA06 and BA45 areas per developmental stage. Serial Xenium sections per sample were treated as technical replicates. Colors indicate significance after Benjamini-Hochberg adjustment for multiple comparisons (p-value < 0.05), log fold changes under absolute value of 0.2 were not plotted. (C-D) Spatial maps of area enriched glutamatergic and GABAergic neurons across infant and adult BA06 and BA45. (E) Spatial maps of *TNC*+ astrocytes in BA06 and BA45 at childhood and adulthood in Xenium 5,000-plex data. Histograms show normalized abundance per cortical depth bin (25 bins). (F) Dotplot showing select differentially expressed genes in Astrocyte *GFAP TNC* between motor (BA06) and Broca’s (BA45) areas in 5,000-plex Xenium data. Sample sizes for 460-plex 10x Xenium ST: n = 4-10 per area per stage Sample sizes for 5,000-plex 10x Xenium ST: n = 1-2 per area per stage.

Our ST analyses validated the area distribution and inferred laminar identity of neuronal subtypes in our consensus cell taxonomy. Glutamatergic neuron subtypes across Brodmann areas mapped to their expected locations, including L2-3 neurons (*L2-3 IT CUX2*), L5 neurons (*L5 IT RORB TRABD2A* and *L5-6NP NPSR1 AS1 ERG*) and L6 neurons (*L6 IT CAR3 THEMIS SMYD1*) subtypes (Figure 4A). We also identified an L4 neuron subtype (*L4 IT RORB OTOGL*) present across areas, despite the agranular nature of premotor cortex (Figure 4A, C). Beyond these, we identified three area-enriched L4-5 glutamatergic neurons. In the adult donor, *L4 IT RORB SLC22A10* and *L5 IT RORB LINC02196* neurons were enriched in Broca’s area, whereas *L5 ET* and *L6 CT ADAMTSL1 SEMA3E* neurons were enriched in motor cortex (Figure 4A-C; FDR=0.05 per donor). These observations are consistent with our snRNA-seq results and established cytoarchitectonic differences between Brodmann areas 44, 45 and 6 (Fig 2C and S7C). Notably, some of these area-specific differences were prominent throughout pediatric development, as they were evident in our infancy, childhood and adolescent samples (Figure 4C). These findings extend previous studies to molecularly define shared and divergent L4/5 glutamatergic neuron subtypes across Broca’s area and premotor cortex.

Furthermore, we uncovered more extensive area enrichment of GABAergic neurons compared to glutamatergic subtypes, consistent with our snMultiome findings. Among neuronal subtypes similarly distributed in cortical layers across both Broca’s and motor areas, *SST CUX2* and *VIP CLSTN2* were positioned in L2-3 and *LAMP5 LHX6* were located in L5-6 (Figure 4A, Figure S11). Beyond these shared profiles, we identified several area- and layer-enriched patterns. The subtypes enriched in Broca’s area included *SST CALB1* interneurons located in upper layers, *SST PROM1* interneurons located in deeper layers and *VIP PTH2R* interneurons without distinct layer localization (Figure 4A-C; FDR=0.05 per donor). *SNCG CADSP2* and *SNCG FOXO1* neurons also showed a trend towards Broca’s area enrichment (Figure S11). In motor cortex, we found enriched *VIP SLC7A11* interneurons located in upper layers (Figure 4A-C). Akin to glutamatergic neurons, some of these patterns (e.g. *SST PROM1* interneurons) were established as early as infancy and childhood (Figure 4B,D).

Finally, we examined spatial distribution of glial cells and identified area differences among astrocytes and oligodendrocytes. Three astrocyte subtypes showed distinct laminar localizations: *HPSE2 GJA1* astrocytes were located in the gray matter, whereas *GFAP GRIA1* and *GFAP TNC* astrocytes were preferentially found in the pial layer/L1 and subcortical white matter (Figure 4E, Figure S12). *TNC+* astrocytes were enriched in Broca’s area compared to pre-motor cortex from childhood onwards (Figure 4E), whereas *GRIA1+* astrocytes were preferentially enriched in pre-motor cortex (Figure S12-20). The *TNC* gene encodes Tenascin-C protein that is crucial for orchestrating neuron and axon migration and facilitating synapses formation^24^. To further assess the area specialization of this astrocyte subtype, we performed differential gene expression analysis of *TNC+* astrocytes from Broca’s versus pre-motor cortex on our 5,000-plex Xenium data. This showed that *TNC+* astrocytes from Broca’s area have enriched expression of glutamate transporters (*SLC1A2*, *SLC1A3*), astrocyte gap junction protein Connexin-43 (encoded by *GJA1*) that underlies astrocyte communication networks^25^, and brain specific angiogenesis inhibitor-1 (encoded by *ADGRB1*) that has been implicated in synaptic pruning by astrocytes^26^. These patterns were detected as early as childhood (Figure 4F, Figure S12), demonstrating molecular specializations of astrocytes that may be involved in supporting the development of neural circuits in Broca’s area. Besides astrocytes, we identified elevated abundance of oligodendrocytes (*ST18 MOBP*) in motor areas compared to Broca’s (Figure S11-12), including a trend towards more prominent gray matter oligodendrocyte populations in the adult pre-motor cortex (Figure S11-12).

Taken together, our spatial transcriptomic analysis resolves shared and divergent cytoarchitectonic features of Broca’s and pre-motor cortex, highlighting prominent regional differences across laminar glutamatergic, GABAergic and astrocyte populations from early pediatric development to adulthood.

### Conservation and divergence of molecular programs in language- and motor-related areas

Functional maturation and regional specialization of the neocortex are accompanied by the progressive establishment of cell type-specific molecular programs, reflected in part by dynamic transcriptional changes. To compare cellular gene expression between Broca’s area and motor cortex and across developmental stages, we conducted differential gene expression analysis using a linear mixed model analysis applied to pseudobulk samples for each cell cluster^27^ (see Methods). This flexible approach enabled us to perform multiple comparisons accounting for both biological and technical sources of variation while conservatively controlling the false positive rate. We first identified changes in gene expression in Broca’s area and motor cortex across developmental stages. Globally, we detected a total of 593 time-dependent differentially expressed genes (DEGs) (FDR < 0.01; Figure 5A and Table S5). These included genes involved in axon guidance, such as the receptors *ROBO1* or *EPHB1,* that are upregulated during infancy in deeper layer glutamatergic neurons, as well as genes associated with synaptic maturation and function, including the neurotransmitter release modulator *SYN3* and the calcium-binding protein *CABP1,* that are upregulated in superficial layer glutamatergic neurons after childhood (Figure S13A,B; Table S6). Conserved molecular programs associated with time-dependent DEGs in GABAergic neurons included pathways related to cell migration and metabolism, consistent with their ongoing postnatal maturation and high metabolic demands^28^. In macroglia, we observed enrichment of terms related to cell-matrix adhesion and regulation of exocytosis, underscoring the roles of glial cells in synaptic and cortical circuit maturation (Figure S13A; Table S6). Approximately two thirds of time-dependent DEGs were found to be specific to either Broca’s area or motor cortex (Figure 5A), and within each class, L2/3 IT neurons were found to be significantly enriched for area-specific time-dependent DEGs (adjusted p-value < 0.05; two- sided Fisher’s exact test), followed by L5 IT and astrocytes (nominal p-value < 0.01; Figure 5B). These subclasses may therefore exhibit more pronounced area-specific molecular signatures in cell developmental trajectories underlying cell maturation and functional specialization across cortical areas.

**Figure 5.**
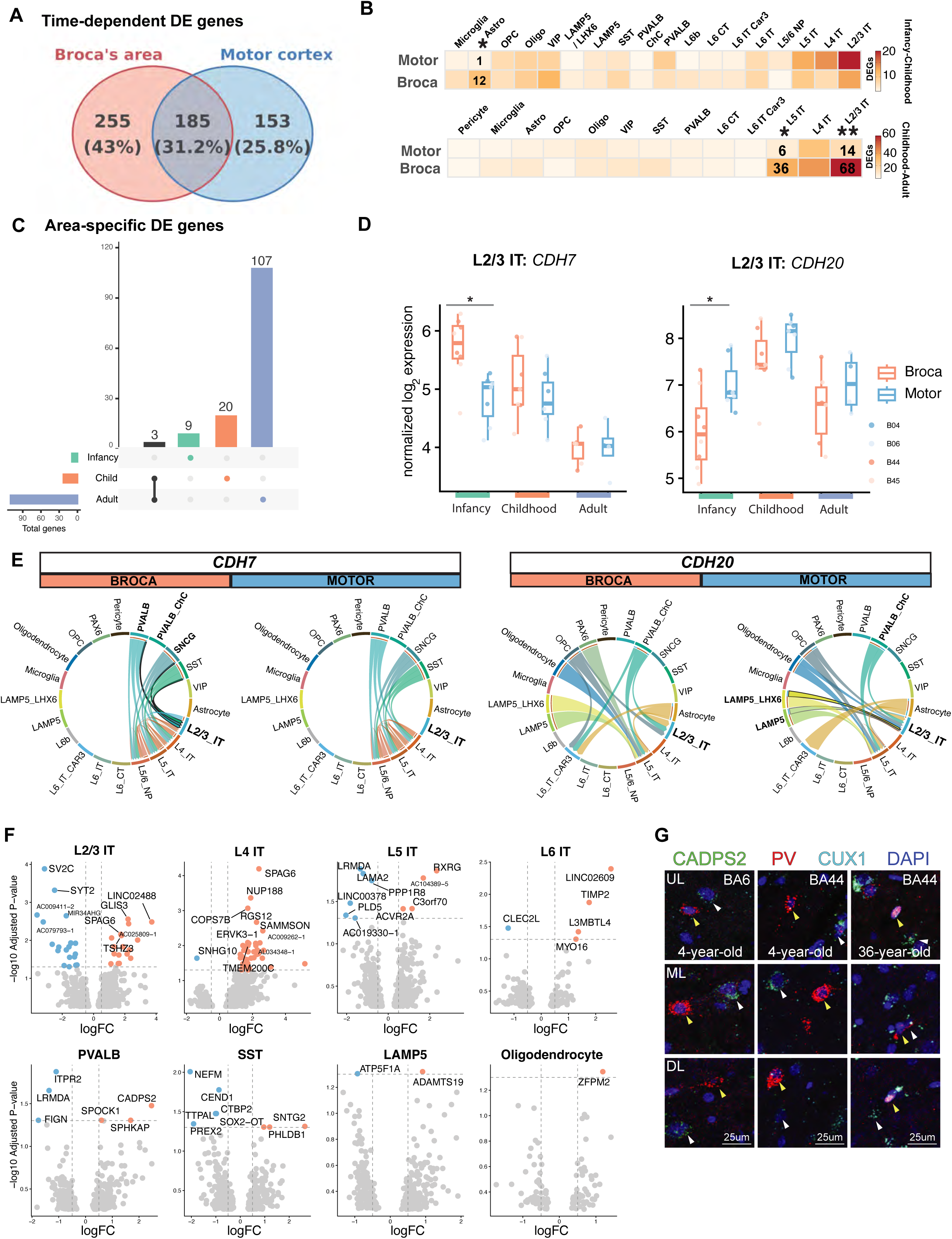
(A) Venn diagram summarizing time-dependent DEGs in either Broca’s area or motor cortex, as well as shared DEGs between these areas. Significant time-dependent DEGs were considered if adjusted p-value below 0.01. (B) Heatmap of enrichment test results for area-specific time-dependent DEGs, highlighting L2/3 IT neurons under adjusted p-value 0.05 (**) and L5 IT neurons and Astrocytes under nominal p-value 0.05 (*) in the transitions infancy to childhood (top), and childhood to adult (bottom). (C) Area-specific DEGs at each developmental stage. Significant results if adjusted p-value below 0.05. (D) Boxplots highlighting differential gene expression of cadherin genes at infancy stage between Broca’s area and motor cortex. (E) Cell-cell communication analysis reflecting incoming signaling towards IT neurons, for homophilic *CDH7* (Broca’s area) and *CDH20* (motor cortex) interactions, highlighting those differentially implicating L2/3 IT neurons with GABAergic neurons. (F) Violin plots summarizing differential gene expression between Broca’s area and motor cortex at adult stage. (G) RNAscope images showing parvalbumin-positive, *CADPS2*-positive PV neurons across areas and selected time points.

**Figure 6.**
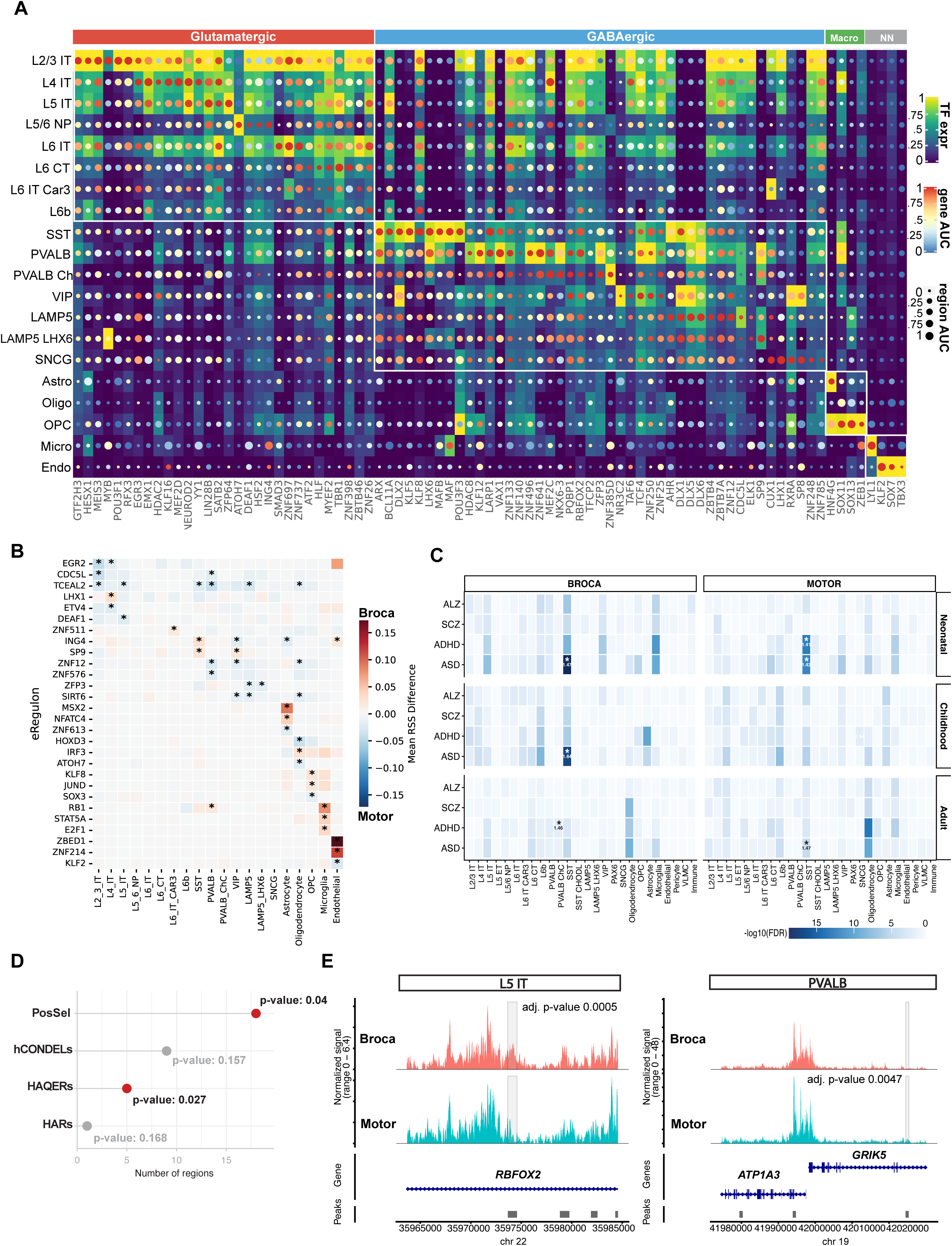
(A) Selected GRNs enriched in different subclasses, summarized by transcription factor expression, region-based AUC scores, and gene-based AUC scores. (B) Selection of GRNs exhibiting differential activity between Broca’s area and Motor cortex per subclass. (C) Area-specific subclass disease relevance score enrichment in ASD, ADHD, SCZ, and ALZ. (D) Permutation-based enrichment test for DARs between Broca’s area and Motor cortex within genomic regions of evolutionary relevance. (E) Evolutionary-relevant DARs within regulatory domains associated with *RBFOX2* and *GRIK5*/*ATP1A3* in L5 IT and PVALB interneurons, respectively.

We then tested for area-specific differential gene expression at each developmental stage. Using our conservative analytic approach, we detected a limited set of DEGs at infancy (n=9) and childhood (n=23) stages, that then cascaded into wider transcriptomic differences in adulthood (n=110 at FDR 0.05; Figure 5C and Table S5). We observed a significant upregulation of the cadherin gene *CDH7* at infancy in L2/3 IT neurons in Broca’s area relative to motor cortex (Figure 5D). *CDH7* expression decayed after infancy to reach levels comparable to that observed in motor cortex (Figure 5D). Concomitantly, we detected an upregulation of cadherin *CDH20* in motor cortex L2/3 IT neurons (Figure 5D). Given their established roles in cell-cell recognition and synaptic specificity^29–32^, the differential expression of cadherins at early postnatal stages, when synaptogenesis is actively sculpting cortical connectivity, suggests that cell type-specific interactions may contribute to the establishment of area-specific neural circuits. To further investigate this possibility, we performed an intercellular communication analysis by modelling ligand-receptor gene expression at cell type resolution^33^. Potential *CDH7*-mediated cell-cell communication was largely restricted to glutamatergic and GABAergic neurons, with interactions between L2/3 IT neurons and PVALB, SST, and SNCG interneurons identified as enriched in Broca’s area (Figure 5E). In contrast, *CDH20*-mediated interactions involved a broader range of subclasses, with interactions between L2/3 IT neurons and LAMP5, LAMP5 LHX6, L5 IT neurons and oligodendrocytes emerging as enriched in motor cortex (Figure 5E; *CDH7* and *CDH20* expression was validated by a combination of RNAscope experiments with selected neuronal markers and spatial transcriptomics, Figure S14A,S15-16). Another differentially expressed gene involved in cell-cell adhesion was the ephrin receptor *EPHA7,* which was significantly upregulated in motor cortex L2/3 IT neurons at infancy (Table S5 and Figure S17). *EPHA7* has been implicated in a neurodevelopmental disorder characterized by intellectual disability and speech delay^34^, and has also been reported to undergo human-specific regulatory control during cortical development^35^. Together, these findings point to an early combinatorial code in which differential expression of cell adhesion molecules may contribute to area-specific neuronal cell-cell communication in Broca’s area.

We identified additional cell type-specific molecular signatures emerging alongside the functional specializations of cortical areas. For example, during childhood, we observed a significant upregulation of the protein tyrosine phosphatase receptor gene, *PTPRK*, in Broca’s area. Notably, *PTPRK* has been associated with educational attainment^36^ and is enriched in the supragranular layers of the inferior frontal gyrus^16^. Our single- cell analysis further localized *PTPRK* differential expression to SNCG interneurons, a population that is preferentially distributed within superficial cortical layers, as confirmed by our spatial transcriptomic profiling (Figure S14B; Table S5). In adults, we identified DEGs involved in signal transduction linked to speech impairment or cognitive function, including *ARHGAP15*^37,38^ and *ARHGEF6*^39^, both downregulated in Broca’s area L2/3 IT neurons, and *FNBP1L*^40,41^ which was upregulated in Broca’s area L2/3 IT and L6 IT Car3 neurons (Table S5). PVALB interneurons, in particular, exhibited molecular signatures indicative of differential intracellular calcium regulation (Figure 5F). In motor cortex, we observed an upregulation of *ITPR2*, a calcium channel in the endoplasmic reticulum^42^ reported to be a candidate susceptibility gene for amyotrophic lateral sclerosis^43^. In contrast, Broca’s area showed increased expression of the calcium-dependent secretion activator protein, *CADPS2* (whose expression in PVALB interneurons was further validated via RNAscope and spatial transcriptomics; Figure 5G and S14C,S18). *Cadps2* knockout mice exhibit selective vulnerability of parvalbumin interneurons and recapitulate several phenotypic features of ASD^29–32^. Moreover, *CADPS2* has been identified as a candidate gene under positive selection in humans and resides within a genomic region significantly depleted of introgressed genetic variants from extinct hominins, a so-called “desert of introgression”^44,45^. Together, these findings highlight candidate genes that may contribute to the molecular specializations of language- and motor-related cortical areas while also conferring cell type-specific disease susceptibility. In the case of *CADPS2*, these observations further raise the possibility that evolutionary pressures unique to the human lineage have shaped molecular programs relevant to both cortical specialization and neurodevelopmental disease vulnerability.

### Subclass- and area-specific gene regulatory activity

Simultaneous profiling of chromatin accessibility and gene expression at single nucleus resolution enables the reconstruction of gene regulatory networks (GRNs). GRNs comprise a collection of interacting transcription factors, target genes, and cis-regulatory elements that collectively establish and maintain cell type-specific gene expression programs, thereby mechanistically informing molecular processes underpinning cell function and susceptibilities to disease. We used SCENIC+^46^ as a computational framework and inferred 427 regulatory networks across subclasses and cortical areas, including 373 governed by TFs acting as activators and 54 as repressors, with a median of 65 genes and 79 unique regulatory regions per regulon (Table S7). Subclass- specific GRN activity was quantified using area under recovery (AUC) scores^46,47^, recovering canonical TFs associated with each subclass (Figure 5A and S19A-B).

These analyses revealed subclass-enriched GRNs controlled by TFs implicated in language- and/or motor- related disorders. Among these, the high-confidence ASD risk gene, *RFX3*^48–50^, emerged as a key regulator in L2-3 IT neurons, while *ARX,* associated with epilepsy-dyskinesia^51–53^ and intellectual disability^54,55^, was enriched in PVALB and SST interneurons (Figure 5A). To assess regulatory divergence between cortical areas, we performed a differential GRN activity test and identified GRNs exhibiting significant differences between Broca’s area and motor cortex (Figure 5B; see Methods). PVALB interneurons and L2/3 IT neurons, as well as microglia and oligodendrocytes showed comparatively higher numbers of subclass-specific differentially active GRNs (Figure S19C). We additionally observed only a modest correspondence between GRNs exhibiting differential activity and differential gene-level expression or peak-level accessibility: only 8 out of 50 differentially active GRNs were enriched in genes showing differential expression, and only 11 out of 50 were enriched in chromatin regions showing differential accessibility (adjusted p-value < 0.05, Fisher’s exact test; Table S7). This could indicate that subclasses across cortical areas differentially leverage the modular architecture of gene regulatory networks, through the cumulative activity of their constituent genes and associated regulatory regions, to implement molecular specializations that extend beyond differential gene expression or chromatin accessibility alone. One example of area-specific regulatory activity is an *MSX2*-driven GRN that exhibited higher activation in Broca’s area astrocytes (Figure 5B). We further found *MSX2* expression was largely restricted to the *GFAP*+*GRIA1+* astrocyte subtype (and some *GFAP+TNC+* cells; Figure S20A-B). This observation mirrors previous observations of *MSX1/2* expression in pial/subpial astrocytes of the macaque neocortex^56,57^. Among predicted target genes of *MSX2* (Table S7), *HS3ST3A1* emerged as a notable candidate. This gene encodes a key enzyme in heparan sulfate biosynthesis, a pathway linked to neurite branching^58^ and synaptic modulation^59^, and exhibited increased expression in Broca’s area in adulthood (in the 10X Multiome data, see Table S5; and validated in the ST data, see Figure S20C). These findings may reflect features of the protracted postnatal maturation of particular astrocyte subtypes and point to area-specific molecular programs driven by differential transcriptional regulatory activity.

### Cell type susceptibility to neurological disorders affecting language and motor function

Language and motor skills are often altered in neurological disorders, and the VLPFC has been associated with conditions such as ASD^60,61^ and dementia^62^. By leveraging our chromatin accessibility atlas at single-nucleus resolution, we tested whether any subclass exhibit increased susceptibility to genetic risk factors associated with ASD, attention-deficit/hyperactivity disorder (ADHD), schizophrenia (SCZ) or Alzheimer’s disease (ALZ) in genome wide association studies (see Methods). We employed SCAVENGE^63^ as a computational algorithm to map GWAS variants to open chromatin regions through a network propagation strategy and to assign cell type specific disease relevance scores (see Methods). We observed disease-specific patterns at the subclass level (Figure S21A), with ASD, ADHD and SCZ genetic risk factors associated with both Glutamatergic and GABAergic neurons (consistent with previous reports^64,65^), while non-neuronal/non-neural cell populations, in particular microglia and astrocytes, exhibited an association to ALZ, consistent with their reported roles in ALZ pathophysiology^66,67^. Analysis of the cellular composition of subclasses with the highest disease relevance scores revealed an increased susceptibility of SST interneurons to ASD and ADHD (Figure 5C). This susceptibility was most pronounced at infancy in both Broca’s area and motor cortex, and persisted into childhood in Broca’s area (Figure 5C). In adulthood, SST and PVALB chandelier interneurons also exhibited associations with ASD and ADHD (Figure 5C). Extending this analysis to our consensus cell taxonomy enabled us to pinpoint particular clusters within each subclass that might drive some of these associations. We found that *SST+PROM1+* and *SST+CUX2+* subtypes exhibited increased susceptibility to ASD/ADHD, especially at early pediatric stages (Figure S21B). Although these findings will require validation in larger single-cell atlases and GWAS cohorts, they are consistent with previous reports of significant molecular changes in SST interneurons in ASD patients^68,69^, and underscore the importance of early pediatric stages for delineating cell type-specific trajectories of susceptibility to neurological disorders.

ASD is frequently associated with language deficits^70^. To more specifically identify implicated cell populations in Broca’s area, we complemented the previous genome-wide analysis with a transcriptomic dataset from BA22 of ASD cases with documented verbal phenotypes^50^ and asked whether transcriptomic signatures distinguishing verbal from non-verbal cases might converge with subclass-specific molecular profiles across language-related cortical networks (comparing DEGs using a per-cell-type sign concordance with an exact binomial test; see Methods). We found that DEGs distinguishing verbal and non-verbal ASD phenotypes showed strong concordance during childhood with area-specific molecular signatures of L4/5 IT neurons (95/126, 75%, p = 4.8 x 10^-9^), L2/3 IT neurons (87/114, 76%, p = 7.6× 10^⁻9^), and PVALB interneurons (60/86, 70%, p = 1.6 × 10⁻⁴; all FDR < 0.01; Figure S22A; Table S8). Specifically, DEGs associated with verbal *versus* non-verbal ASD phenotypes in L2/3 IT neurons were enriched among genes down-regulated in Broca’s area, whereas those in L4/5 IT neurons and PVALB interneurons were enriched among genes up-regulated in Broca’s area relative to motor cortex (Figure S22B). No significant concordance was observed in adulthood. These findings suggest that transcriptional alterations associated with verbal phenotypes in ASD preferentially converge on molecular programs of specific excitatory and inhibitory neuronal subclasses across language-related cortical networks during early postnatal development.

### Genomic specializations of ventral prefrontal cortex regulatory regions shed light on the evolution of linguistic ability

We interrogated evolutionary specializations of genomic regions that might have contributed to the emergence of linguistic abilities and fine motor skills characteristic of our species. We tested whether differentially accessible regions (DARs; adj. p-value < 0.01, Wilcoxon rank sum test; Table S9) between Broca’s area and motor cortex are over-represented within genomic regions that have diverged during human evolution. These genomic specializations include human-specific deletions in conserved elements (hCondels)^71^, human accelerated regions (HARs)^72^, human ancestor quickly evolved regions (HAQERs)^73^, and human positively selected regions^74^. Consistent with previous reports linking HAQERs to spoken language^75^, we observed a significant enrichment of DARs within HAQERs (permutation test; p-value 0.03; Figure 5), noncoding genomic regions that underwent accelerated evolution after the divergence of the human and chimpanzee lineages approximately six million years ago but remained constrained across hominins^73^. In addition, we observed a significant enrichment within positively selected genomic regions that differentiate present-day humans from extinct hominins for which high-quality genomes are available^45^ (permutation test; p-value 0.04; Figure 5D). These findings highlight the acquisition of putative regulatory genetic variants over the past one million years that may have also contributed to the emergence of linguistic abilities in the *Homo sapiens* lineage. We next annotated regulatory domains^76,77^ associated with these significant DARs and identified genes under putative differential regulation between Broca’s area and motor cortex at each developmental stage (Table S10). Several prominent candidate genes emerged during early pediatric stages, including *RBFOX2,* associated with a DAR in L5 IT neurons during childhood, that is a regulator of alternative splicing and neuronal maturation^78,79^ previously associated with language abilities^80^ and variation in grey matter thickness in language-related cortical areas^81^ (Figure 5E). We also identified a DAR in PVALB interneurons during childhood within a regulatory domain encompassing *GRIK5* and *ATP1A3. ATP1A3* encodes a Na+/K+ ATPase pump and is mutated in several neurodevelopmental disorders including bilateral perisylvian polymicrogyria^82^, childhood hemiplegia with language impairments^83,84^, and childhood-onset schizophrenia^83^ (Figure 5E). Although the inherent sparsity of ATAC data provides only a limited view of cellular chromatin profiles, these analyses uncover a set of differentially accessible regions between Broca’s area and motor cortex that might have contributed to cell type-specific molecular specializations related to the emergence of linguistic and fine motor abilities.

## DISCUSSION

We establish a transcriptomic, epigenomic and spatial atlas of Broca’s area and the adjacent motor cortex during pediatric stages of brain development. Our multi-layered developmental analysis across biological scales and extending to the adult human brain reveals cortical area-specific dynamics in gene expression and regulatory programs, cellular composition and cytoarchitectonic specializations within core cortical areas underlying language and motor function.

### Molecular and cellular correlates of language- and motor-related developmental milestones

Language learning in children follows a stereotyped sequence of developmental milestones that typically culminates in the acquisition of language, raising the question of whether distinct molecular and cellular programs underlie these developmental milestones^85–87^. At the molecular level, we identified a transient differential expression of the cadherin genes, *CDH7* and *CDH20*, as well as the ephrin receptor gene, *EPHA7*, distinguishing superficial layer neurons of Broca’s area from those of the adjacent motor cortex during infancy. These differences may contribute to area-specific cell-cell communication, consistent with ligand-receptor analyses predicting distinct interactions between superficial layer neurons and GABAergic and nonneuronal subclasses. Consistent with this hypothesis, cell adhesion molecules play a prominent role in developmental mechanisms of human cortical arealization and corticocortical connectivity^88^, and previous studies have established instructive roles for cadherins in synaptic specificity and neural circuit assembly in the rodent retina^30^, hippocampus^31^ and cerebral cortex^32^. Such an early combinatorial adhesion code could therefore enable spatiotemporally precise target recognition and integration of interneurons and glial cells, thereby supporting the assembly and function of area-specific neural circuits. Alterations in the expression levels of these molecules could consequently perturb normal developmental trajectories. Although *CDH7* and *CDH20* have not, to our knowledge, been directly implicated in language or motor disorders, mutations in *EPHA7* have been associated with a range of neurodevelopmental phenotypes including absent speech and developmental delay^34^. Intriguingly, dynamic regulation of cadherin 7 expression, as found here in Broca’s area, has also been observed in vocal motor centers in songbirds, specifically the Bengalese finch where cadherin 7 is expressed in the robust nucleus of the arcopallium during the transition from the sensory to the sensorimotor vocal learning stage^89,90^. Moreover, disruption of cadherin 7 expression markedly impairs song learning in this species^89,90^. This suggests that cadherin-mediated mechanisms may play a conserved evolutionary role in aspects of vocal communication^91^.

*CDH7*-mediated intercellular communication, enriched in Broca’s area, is predicted to occur between L2/3 IT neurons and PVALB interneurons. Intriguingly, PVALB interneurons in adult Broca’s area significantly upregulated *CADPS2*, a calcium-dependent regulator of exocytosis that bears strong signatures of positive selection in the human lineage, yet whose functional contribution to human brain evolution remains unresolved^45,92,93^. Our findings nominate differential *CADPS2* expression in PVALB interneurons as a candidate molecular specialization of the VLPFC with potential evolutionary relevance to language-related neural circuitry. Enriched *Cadps2* expression has likewise been reported in songbird brain regions involved in song learning^94^, reinforcing the idea that species with vocal learning abilities may share conserved molecular specializations^95–97^. Future comparative studies across primates and other species with vocal learning abilities may further reveal both shared and divergent features underlying the neural implementation of language and communication.

At the cell population level, our findings highlight pronounced spatiotemporal dynamics of glia populations across the cortical areas examined here, coinciding with time-dependent gene expression changes related to cell matrix organization and synaptic regulation. These observations further support a role for glia in the development and maturation of neural circuits^98,99^. Collectively, we captured significant changes in molecular and cellular axes of variation related to synaptic connectivity within the ventral prefrontal cortex during early pediatric stages. These changes might significantly contribute to multiscale developmental programs that support the progression of developmental milestones required for the proper acquisition of language and motor skills.

### Cytoarchitectonic specializations of Broca’s area and motor cortex

By combining Xenium spatial transcriptomics with serial sectioning of time- and area-matched samples co-profiled by 10x Genomics Multiome, we spatially mapped cellular and molecular axes of variation onto the cytoarchitectonic landmarks of Broca’s area *versus* motor cortex. Our spatial reconstruction of cortical area cytoarchitecture at single-cell resolution orthogonally validated and extended our snMultiome findings, identifying clear area enrichment of multiple excitatory and inhibitory neuron subtypes as well as astrocytes. Across developmental time, we observe changes in cell type population sizes that may be partly explained by cell death^100^ as well as by shifts in combinatorial marker gene expression that may accompany postnatal cell type maturation. Between cortical areas, while the arealization of L3-5 neurons has been documented^12^, our analysis molecularly defines shared and divergent sets of excitatory neurons across Broca’s area and adjacent motor cortex, including several L2-5 IT neuron subtypes enriched in the former that may underlie the connectivity of Broca’s area to other cortical and subcortical regions. Furthermore, we found extensive regional differences across interneurons, spanning both upper and lower layer subtypes, indicating that area patterning of inhibitory neurons plays a prominent role in cortical specialization across language and motor circuits. Notably, these specializations extend to laminar astrocytes enriched in Broca’s area that express genes related to excitatory neurotransmission, synaptic pruning and multi-cellular astrocyte networks. We find that these neuronal and glial specializations are prominent from early postnatal development, suggesting they are established by early developmental area patterning mechanisms. Together, our findings indicate that, over the prolonged postnatal development of the human brain, extending beyond adolescence, cortical regions supporting language and fine motor control undergo progressive area- and stage-specific specialization, reflected in the emergence of cell type-specific gene expression programs, patterns of cell-cell communication, and laminar organization.

### Evolutionary specializations and vulnerability to disease implicating language and motor function

The ventrolateral prefrontal cortex has significantly expanded over the course of human evolution^101^, and elucidating its evolutionary and developmental trajectory may provide insights into susceptibilities to neurodevelopmental and neuropsychiatric disorders. Our evolutionary-informed epigenomic investigation of Broca’s area and motor cortex yields two main insights, albeit with the caveat that the inherent sparsity of ATAC data precludes deep phenotyping of cellular chromatin landscapes. First, differentially accessible regions that spatiotemporally delineate the cellular epigenomic profiles of Broca’s area and adjacent motor cortex are enriched in genomic specializations that distinguish our species from our closest living primate relatives and archaic hominins. The temporal resolution afforded by the joint investigation of these genomic specializations invites the suggestion that genetic changes associated with phenotypic adaptions relevant to human linguistic abilities accumulated progressively over the course of million years in *Homo sapiens*, rather than in a single abrupt evolutionary event. Future work integrating epigenomic and genomic data across extant and extinct species may further illuminate the rate at which derived phenotypic traits relevant to human cognition evolved. Second, some of these genomic specializations may be associated with the regulatory control of genes that confer increased susceptibility to neurological disorders. This is illustrated perhaps most prominently by *ATP1A3*, a gene whose mutations carry profound clinical consequences^82^. We further uncovered cell type-specific patterns of vulnerabilities to disease, prominently somatostatin-positive interneuron subtypes that exhibit a pronounced susceptibility to ASD/ADHD during early developmental stages. These findings extend prior reports^68,69^ and prompt the question of how cellular susceptibility to neurodevelopmental and neuropsychiatric conditions evolves over developmental time. Finely resolved maps of the dynamic molecular and cellular landscapes spanning prenatal and early pediatric development will be critical for understanding how developmental cell trajectories contribute to disease susceptibility and variability in penetrance.

### Limitations

Primary tissue samples across early postnatal developmental stages remain limited. Increasing sample size will be required to more deeply phenotype the molecular and cellular dynamics of early pediatric brain development and resolve the progressive refinement of neural circuits into adulthood. Adolescent samples remain under- represented in our datasets, and future studies incorporating larger cohorts will substantially increase the power of single-cell analyses, particularly in light of the challenges associated with estimating cell type proportions from single-cell data and the sparsity of single-cell ATAC-seq chromatin accessibility data.

## METHODS

### Primary tissue samples

Human brain tissue samples were obtained from two independent sources (Table 1). Four de-identified samples (9 tissue samples), representing infancy, childhood, adolescence and adult developmental ages were provided by the UCSF Pediatric Neuropathology Research Laboratory (PNRL, PI: E.J.H) and the Autopsy Service in the Department of Pathology at the University of California San Francisco. All samples were collected with previous patient consent in strict observance of the legal and institutional ethical regulations. Autopsy consent and all protocols were approved by the Human Gamete, Embryo, and Stem Cell Research Committee and the Institutional Review Board (IRB) at the University of California San Francisco. All cases received extensive neuropathological evaluations to rule out the presence of neurological diseases. Subjects from both genders were used for this study. Information regarding age and gender is summarized in Figure 1A. Briefly, fresh brain tissues from the Broca’s Area (Brodmann Area 44 and Area 45) and premotor cortex (Brodmann Area 6) and motor cortex (Brodmann Area 4) were dissected and serially sectioned using razor blades with alternate sections snap-frozen or fixed in 4% paraformaldehyde for further analysis. Tissue samples were stored at -80°C for long-term preservation. These samples were used for both single-nucleus Multiome sequencing, histology, spatial transcriptomics analyses and longitudinal RNAscope assays. An additional 32 tissue samples from 8 donors, spanning infancy, childhood and adult stages were obtained from the University of Maryland Brain and Tissue Bank through the NIH NeuroBioBank (NBB). These included cortical regions BA4 (premotor), BA6 (motor), BA44 (Broca’s area), and BA45 (Broca’s area). The acquisition and experimental use of these tissues were conducted under UCSF IRB approval, in accordance with all relevant ethical guidelines. A comprehensive list of the samples used for single-nucleus Multiome and spatial transcriptomics analyses is provided in Table 1.

### Nissl Staining

Tissue samples obtained from the UCSF PNRL were fixed in 4% paraformaldehyde (PFA) in PBS and sectioned at 16um thickness onto SuperFrost Plus glass slides. The sections were air-dried and baked at 60°C to ensure adherence. Slides were then immersed overnight in a 1:1 alcohol-chloroform solution and rehydrated through a graded ethanol series (100% and 95%) to distilled water. Sections were stained in prewarmed 0.1% cresyl violet solution for 5-10min, to enhance visualization of neuronal cytoarchitecture. After rinsing in distilled water, slides were differentiated in 95% Ethanol for 15min, followed by dehydration in 100% ethanol for 10min. The sections were then cleared in xylene for two 5min incubations and mounted using a xylene- compatible mounting medium. Stained sections were imaged under a bright-field microscope to assess the overall neuronal morphology and cortical organization.

### Nuclei isolation and single-nucleus multiomics data generation

A detailed nuclei isolation protocol has been previously described (dx.doi.org/10.17504/protocols.io.eq2lyj3nplx9/v1). Briefly, all fresh frozen tissue samples (50-100mg), maintained on ice during processing, were homogenized using a pre-chilled 7ml Dounce homogenizer, containing 1.5ml cold homogenization buffer (HB- 20 mM Tricine-KOH pH 7.8, 250 mM sucrose, 25 mM KCl, 5 mM MgCl_2_, 1 mM dithiothreitol, 0.5 mM spermidine, 0.5 mM spermine, 0.3% NP-40, 1× cOmplete protease inhibitor (Roche), and 0.6 U ml^−1^ RiboLock (Thermo Fisher Scientific). Tissues were homogenized 12 strokes of the loose pestle and 15 strokes with the tight pestle. The resulting lysate was centrifuged at 500g for 5min at 4°C to pellet nuclei. The pellet was resuspended in an iodixanol density gradient (25%, 30%, and 40%) and centrifuged at 3,000g for 20min (acceleration set to 0, deceleration set to 1). Clean nuclei were collected at the 30-40% interface and diluted in wash buffer containing 10 mM Tris-HCl pH 7.4, 10 mM NaCl, 3 mM MgCl_2_, 1 mM dithiothreitol, 1% BSA, 0.1% Tween-20, and 0.6 U ml^−1^ RiboLock (Thermo Fisher Scientific). Nuclei were pelleted again by centrifuging at 500g for 5min, at 4°C and resuspended in 10x Genomics diluted nuclei buffer. Nuclei were counted manually using a hemocytometer, adjusted to a final concentration of 3,220 nuclei per ul, and processed following the 10x Genomics Chromium Next GEM Single Cell Multiome ATAC + Gene Expression Reagent Kits following the manufacturer’s guidelines. We targeted 10,000- 12,000 nuclei per sample per reaction. Libraries were multiplexed and sequenced on the Illumina NovaSeq 6000 and NovaSeq X sequencing platforms, targeting an average depth of 25,000 paired-ended reads per nucleus for both ATAC and gene expression data.

### Single-nuclei multiome data preprocessing

Cell Ranger ARC analysis pipelines (v2.0.2; 10X Genomics) were used to process 10X Genomics single-nucleus multiome (RNA+ATAC) sequencing raw data. 10X Genomics human reference packages GRCh38 and GENCODE v32/Ensembl98 were used for read alignment and genome annotations. High-quality cells were retained according to the following cell-based metrics, applied per sample: A maximum of five times the median absolute deviation of unique transcripts and number of detected genes, with at least 400 genes detected; less than one percent of total reads assigned to mitochondrial reads; total ATAC fragment counts between 400 to 100,000; transcription start site enrichment score greater than one; ratio of mononucleosome to nucleosome-free fragments below two. As global filters, cells with number of detected genes or counts above the respective 95th percentile values were also filtered out to further exclude potential outlier cells. Ambient RNA was estimated using SoupX^102^ and homotypic and heterotypic doublet detection was carried out using scDblFinder^103^ and Amulet^104^.

### Hierarchical iterative clustering

Subclass level annotation was performed based on well-established markers as well as integration of our data with reference datasets^105,106^. Clusters were defined in the adult samples using the iterative clustering package hicat^107^ by pooling cells from all four cortical regions. First, mitochondrial, ribosomal, sex-specific genes, and genes detected in less than four cells were removed from the expression matrix. Cells were grouped into neighborhoods based on their subclass assignment and hicat clustering was run on each neighborhood separately. Every iteration of hicat clustering consisted of high-variance gene selection, dimensionality reduction, and Jaccard-Louvain clustering applied recursively until differential expression between clusters no longer passed a threshold differential expression score (sum of -log10 of adjusted p-value over significantly differentially expressed genes). This process was run 50 times on 80% subsamples, then consensus clusters were consolidated based on how often cells clustered together across iterations. Consensus clusters were inspected for their distribution across individuals to identify clusters dominated by individual batch effects. hicat clustering was run once again for 100 iterations by passing the first principal component of batch-specific gene expression. Remaining batch effect clusters were either merged with the nearest cluster or removed (in the case of clusters with low-quality cells) until all clusters were reasonably distributed across individuals. High-confident clusters were then annotated by differentially expressed marker genes relative to other clusters globally, clusters in the same neighborhood, or clusters in the same subclass. Finally, following previous work^108^, cluster annotations were then sequentially transferred to earlier time points using reciprocal PCA-based label transfer. Donors were well represented across clusters and developmental stages; however, a small number of clusters in early developmental stages (4 childhood and 4 infancy) were not represented by at least one individual donor. A few clusters, likely under-sampled in our dataset, were not detected in at least one Brodmann area: *PAX6 PALMD* at childhood, and *PAX6 PALMD* and L5-6NP *NPSR1-AS1 PCED1B* at infancy.

### Cell compositional analysis

We evaluated cell proportion changes across developmental stages and between cortical areas by leveraging a case-control compositional data analysis framework^21^. This consists of an isometric log-ratio transformation of cell type proportions per sample and an estimation of statistical significance against an empirical distribution. This approach is robust to variability between samples in single-cell data and accounts for dependencies between cell types in proportion tests. We performed an independent analysis of compositional changes at Subclass and taxonomy cluster levels, and in a pairwise-manner comparing infancy and childhood stages, and Childhood and Adult stages, in Broca’s area and motor cortex independently. We also directly compared these areas at each developmental stage. To avoid potential compositional shifts caused by cell clusters with low representation in the dataset, we required at least a minimum of 10 cells per comparison group.

### Intercellular communication analysis

Intercellular communication modelling using the gene expression modality was implemented using CellChat (v2.1.2)^33^. To test whether differential gene cadherin expression impact cell-cell communication, we considered infancy stage samples only. Additionally, we augmented the default CellChat database to include a repertoire of adhesion-mediated interactions using high-confidence cadherin interactions derived from STRING v12.0 (score >0.9)^109^. For each dataset, ligand–receptor interactions and communication probabilities were inferred at the subclass level using default parameters. Interactions involving sender or receiver populations represented by fewer than ten cells were excluded. To compare intercellular communication between Broca’s area and motor cortex, region-specific CellChat objects were merged and differential interaction analyses were performed using CellChat’s comparative framework. Differences in the number and strength of inferred interactions, signaling pathways, and ligand–receptor pairs were assessed using permutation-based statistical procedures implemented in CellChat. Weighted directed communication networks capturing cadherin-mediated signaling were visualized using chord diagrams, with a particular focus on incoming and outgoing signaling involving L2/3 IT neurons and the strength of communication between subclass pairs.

### Gene regulatory networks inference

Reconstruction of GRNs was performed within the SCENIC+ framework^46,47^ leveraging both RNA- and ATAC- sequencing data modalities. Due to inherent noise and sparsity of 10X snMultiome data, and to overcome the computational limitations of processing large-scale datasets within the SCENIC+ environment, we performed GRN reconstruction following a pseudobulk strategy. Briefly, single cells were aggregated into metacells using SEACells^110^ applied independently to each sample. Samples with fewer than 2,000 cells were excluded from the analysis and the number of metacells per sample was set to approximately 1 per 75 cells. Archetype initialization was performed using 10 eigenvalues and SEACells model was fit with a minimum of 10 and maximum of 100 iterations and default convergence epsilon 1e-5. Metacells were summarized using soft assignments and cell-metacell pairs were retained if a minimum model weight of 0.05. Metacell quality was assessed by computing subclass purity (proportion of dominant cell subclass within each metacell), compactness (variance among constituent cells) and separation (distance to the cells from nearest neighboring metacell). An aggregated raw count matrix across samples was generated using SEACell function *genescores.prepare_multiome_anndat*.

For GRN reconstruction and following SCENIC+ guidelines, topic modelling was performed though the Latent Dirichlet Allocation algorithm with Mallet parallelization, with evaluation of optimal number of topics based on four different metrics available in SCENIC+ framework. For motif analysis, topic modelling was performed by applying the Otsu method for topic binarization, with the selection of top 3000 regions per topic; additionally, differentially accessible regions were identified using a Wilcoxon rank sum test requiring at least a log fold- change higher than 0.5 and adjusted P-value below 0.05. Next, the motif enrichment analysis implemented in the pycisTarget suite was used to identify significant associations between transcription factors and candidate regulatory regions. Finally, transcription factor-to-gene expression as well as candidate regulatory regions-to- gene expression associations were inferred using default parameters on the Gradient Boosting Machine regression implemented within the SCENIC+ framework. Pearson correlation determined positive or negative relationships for the associations. After filtering for a minimum of ten genes and extended annotations only if no direct annotations available, we obtained a total of 427 enhancer driven regulons. To evaluate subclass- specific activity of GRNs identified, regulons specificity scores (RSSs) were computed using default SCENIC+ algorithm based on Jensen-Shannon divergence^111^ independently for each subclass. Next, to estimate differential GRN activity between Broca’s area and Motor cortex at the subclass level while mitigating biases arising from unequal subclass size (cell number) between areas, we implemented a bootstrapping procedure. Pairwise comparisons of subclass RSSs were performed after randomly down sampling the larger subclass in either Broca’s area or Motor cortex to match the size of the smaller subclass. Bootstrap resampling with replacement (1,000 iterations) was then performed to estimate mean RSS difference and 95% confidence intervals. Differential regulon activity was considered significant if a) confidence intervals excluded zero, b) estimated RSS difference exceeded a global effect size threshold defined as the mean plus three standard deviations from the absolute RSS difference distribution across all regulons, and c) TF governing the regulon is sufficiently expressed (normalized expression > 0.5) in the relevant subclass.

To test whether GRNs exhibiting differential activity were also enriched for genes/peaks showing differential expression/accessibility, we identified DEGs and DARs using more lenient thresholds (adjusted p-value < 0.05, no minimum log fold-change cutoff) than those used elsewhere, to account for the conservative nature of our analytical approach (see below for differential expression/accessibility analyses) and to capture subtler molecular divergences in regulatory programs that a strict DEG/DAR thresholding might fail to detect. For each modality, we tested subclass-specific DEGs/DARs against subclass-specific regulon target gene/region sets by constructing contingency tables and applying a one-sided Fisher’s exact test to assess over-representation. Resulting p-values were corrected for multiple testing across all subclass-by-regulon tests within each modality using the Benjamini-Hochberg procedure, and enrichments with an FDR-adjusted p-value < 0.05 were considered significant.

### Differential gene expression analysis

A pseudobulk-based, linear mixed model analysis^27^ was implemented to identify differentially expressed genes in each cell cluster across developmental stages and cortical areas. Quantification of relevant sources of variation in gene expression was performed using the *fitVarPart* function in the dreamlet package^27^ and differential expression test was implemented using *dreamlet* function. Contrasts were designed to compare cell cluster gene expression between Broca’s area and motor cortex and age groups, accounting for other sources of variation (in particular, our sample metatadata categories: Individual, Post-mortem time interval, Sex and Laboratory). Subclasses were considered for analysis only if at least 15 cells were retained in each experimental condition (consisting of Brodmann area and developmental stage). Significant differential gene expression was considered if adjusted p-value < .05 (Benjamini-Hochberg correction) unless otherwise noted. Gene Ontology (GO) enrichment analysis was performed using the ViSEAGO R package^112^ (v1.18.0): To assign DEG to biological processes and compare across cortical areas, for each cell class, genes were partitioned into significant (adj. p-value < 0.01) and background sets independently for Broca’s area and motor cortex. Gene to biological process (BP) assignment was tested separately per area using Fisher’s exact test with the classic algorithm implemented in topGO^113^. Enriched terms from both regions were merged at a significance threshold of p-value < 0.05. Semantic similarity between enriched GO terms was computed using the Wang distance metric, and GO terms were clustered hierarchically using Ward’s D2 agglomeration on the resulting similarity matrix.

### Xenium Spatial transcriptomics

Brain tissue was fixed with 4% PFA, embedded in OCT and frozen in isopentane-dry ice slurry. Tissue blocks were serially sectioned at 10 μm thickness and a section of each cortical area per age was collected on the same xenium slide in order to have representation of all three cortical areas per age for batch effects. All sections were collected on cold xenium slides maintained in the cryostat, before moving them for storage at –80°C. The first section of each series was used for Xenium Prime (5K panel) profiling, with the next sections profiled with the custom panel. Sections for Xenium v1 (custom panel) were collected at 120 μm intervals in order to profile a 400 μm thick tissue block at 4 different levels.

All sections were processed following the modified 10x Genomics guide for fixed-frozen tissue (https://kb.10xgenomics.com/s/article/17968908868877-Are-fixed-tissues-embedded-in-OCT-compatible-with-Xenium). Briefly, sections were rehydrated through a graded ethanol series (user guide CG000662) and de-crosslinked (CG000578) before proceeding with probe hybridization, ligation, amplification, and cell segmentation staining, following either the Xenium In Situ Gene Expression with Cell Segmentation Staining User Guide (CG000749) for the custom panel, or Xenium Prime In Situ Gene Expression with Cell Segmentation Staining User Guide (CG000760) for the Prime 5K Human Pan Tissue and Pathways Assay panel.

### Label Transfer using rDOT

The AnnData object of 10X multiome was subset into separated objects by Brodmann area and developmental stage. The raw count matrix of each object is subsequently normalized to the target sum of 1 x 10^4^ and log transformed. Highly variable genes were computed using the flavor of ‘seurat_v3’^114^. The final AnnData object was subset by the highly variable genes and used as reference for label transfer. The R package ‘DOTr^115^’ was performed for label transfer at matching Brodmann area and developmental stage. The raw count matrix of spatial data and spatial coordination of each cell was retrieved as input. Cell type reference was generated using the raw count matrix, highly variable genes and the 69 cell type labels with a parameter of reference subcluster size at 3. A DOT object was created with above information and performed high resolution cell type label transfer with ratios weight at 0. The output DOT plotting file containing predicted cell types was merged with the ST dataset by cell index.

### Cortical Layer and cortical depth assignment

The 10X Xenium dataset were integrated using Harmony^116^. Coarse clustering was first used to identify and annotate white matter, grey matter and meninges. For cortical layer annotation, gene scores were calculated for marker genes associated with cortical layers. Marker scores were then used to assign cortical layers per cell. Cells localized within a layer expressing layer-specific marker signatures were annotated accordingly. Cells that could not be confidently assigned based on marker scores alone were subsequently annotated using their spatial proximity to neighboring clusters with confident cortical layer assignments. To further validate, we subsequently performed spatial-aware clustering BANKSY^117^ using scaled gaussian model with resolution of 0.20 and k number at 40.

To define cortical depth from pia-vasculature to white matter, a polygon region of interest was manually delineated on each tissue section in spatial coordinates. The polygon boundary was annotated such that edges facing the meningeal surface were designated as pia boundaries, and edges facing the subcortical white matter were designated as white matter (WM) boundaries, classified by their geometric proximity to the two endpoints of a manually placed reference axis (p1 at the pia, p2 at the WM surface). Transition edges connecting the two boundary types were treated as lateral walls.

For each cell within the polygon, a normalized depth coordinate *t* ∈ [0, 1] was computed as the distance-ratio:

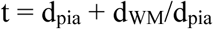

where *d*_pia and *d*_WM are the minimum perpendicular Euclidean distances from the cell’s spatial coordinates to the nearest pia boundary segment and the nearest WM boundary segment of the polygon, respectively, with the foot of the perpendicular clamped to the segment endpoints. Cells falling outside the polygon were excluded. This yields *t* = 0 at the pia surface and *t* = 1 at the WM boundary.

A reference point marking the grey-white matter junction was manually placed on the reference axis, and its depth coordinate *t*_ref was computed by the same distance-ratio method. Depth was then rescaled to a normalized coordinate:

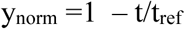

such that *y*_norm = 1 at the pia, *y*_norm = 0 at the grey-white matter boundary, and *y*_norm < 0 for cells extending into white matter. Cells were binned along *y*_norm into equal-width bins spanning from the pia to the reference depth. Cell abundances of cell types were normalized by the surface area of each bin.

### Differential abundance analysis

The AnnData object of 10X Xenium ST dataset was subset by age. For each developmental stage, we performed scCODA^118^ at cell level with the 69 Cell Types label as cell type identifier. For each developmental stage, we compared the cell type abundances between cortical areas (Broca’s and Motor). The Immune *SKAP1 CD44* was used as reference cell type. The cell type abundances were evaluated with FDR < 0.05.

### Integration of genetic risk variants from GWAS with chromatin accessibility profiles at single-cell resolution

We used SCAVENGE^119^ to integrate GWAS variants with our single-nucleus chromatin accessibility atlas. We considered four neuropsychiatric disorders: ASD, ADHD, SCZ, and Alzheimer’s disease, using GWAS summary statistics available from https://figshare.com/articles/dataset/asd2019/14671989 (ASD), https://figshare.com/articles/dataset/adhd2022/22564390 (ADHD);https://figshare.com/articles/dataset/cdg2018-bip-scz/14672019 (SCZ) and https://vu.data.surfsara.nl/index.php/s/jVlyt1m9Bb2mAki/download?path=%2F&files=PGCALZ2sumstatsExcluding23andMe.txt.gz (ALZ).

Briefly, following SCAVENGE standard workflow, ABF fine-mapping (prior = 1×10⁻⁴) was performed on 1 Mb windows centered on GCTA-COJO–derived sentinel variants^120,121^. Variants were ranked by posterior probability to define 95% credible sets and fine-mapped posterior probabilities were imported into gchromVAR to compute trait-relevant scores (TRS) for each cell. To minimize batch effects, we used the batch-corrected LSI embedding (top 40 dimensions) for nearest-neighbor graph construction (k = 30 mutual nearest neighbors) and subsequent network propagation (γ = 0.05). Seed cells were defined as those in the top 5% of gchromVAR z-scores, and TRS values were scaled using a scale factor derived from the top 1% of z-scores, followed by outlier capping at the 95th quantile and min-max normalization. Cells in the top 0.1% of TRS scores were considered trait-relevant, with statistical significance assessed through 1,000 permutations (P < 0.001). Cell type enrichment was evaluated using two-sided Fisher’s exact tests with Benjamini–Hochberg correction; subclasses or subtypes with FDR < 0.05 and odds ratio > 1.4 were considered significantly enriched for trait-associated variants.

### Cross-area sign-concordance analysis

Single-nucleus RNA-sequencing profiles of BA22 were obtained from post-mortem tissue of ASD donors with documented verbal/non-verbal phenotypes, originally annotated into 38 neuronal and non-neuronal cell subtypes^50^. BA22 fine subtypes were mapped to coarser labels in our study by canonical subclass identity (e.g. L2_3_IT1–4 to L2/3_IT; PValb_1–4 to PVALB, or L4_5_IT1-3 to L4/5 IT), yielding a direct one-to-one correspondence between studies. Within ASD male donors over two years of age (30 verbal, 11 non-verbal), differential expression between verbal and non-verbal cases was computed per cell type with FindMarkers (wilcoxon rank-sum test, pseudocount.use=0), using the default gene-level pre-filters of logfc.threshold = 0.1 and min.pct = 0.01; genes passing these filters were reported with an average log fold change and p value. These were intersected per matched cell type with Broca vs motor cortex differential expression across infancy, childhood and adulthood (L4/5 IT matched the pooled Broca vs motor cortex L4_IT and L5_IT sets). For each cell-type × age panel, candidate DEGs were defined by p < 0.05 in both contrasts, and genes with non-finite log fold changes were excluded. Under the null of independence, sign agreement between BA22 average log fold change and Broca’s area vs motor cortex. log fold change follows Binomial(n_overlap, 0.5). We tested for excess concordance with a one-sided exact binomial test (binom.test, alternative = “greater”), correcting across all panels by Benjamini–Hochberg FDR.

### Enrichment of differentially accessible regulatory elements in evolutionary relevant genomic regions

We asked whether regulatory elements differentially accessible between Broca’s area and motor cortex are enriched in genomic regions of relevance for human evolution; specifically, human accelerated regions^72^, human ancestor quickly evolved regions^73^, human-specific deletions in conserved regions^71^, and putative positively selected genomic regions^74^. Differentially accessible regions were identified by performing a Wilcoxon rank-sum test^122^ at each developmental stage and significance was considered if adjusted p-value was below 0.01 and log fold change higher than 1.25. We used regioneR^123^ package to compute the overlap between our query and evolutionary-relevant genomic regions and to perform permutation tests to quantify significance of enrichment, controlling for size of and distance between regions.

### Immunohistochemistry

Frozen tissue was sectioned in 16um thickness onto Fisherbrand^TM^ SuperFrost glass slides. Antigen retrieval was performed using 1X boiling Citric acid-based unmasking solution (Vector Laboratories) for 15min. Sections were blocked for 1h at RT in Donkey blocking buffer (DBB) containing 5% donkey serum, 2% gelatin and 0.1% Triton X-100 in PBS. Primary antibodies were diluted in DBB and incubated overnight at 4°C, followed by three washes with 0.1% PBST (PBS+Triton X-100). Sections were then incubated with AlexaFluor-conjugated secondary antibodies (ThermoFisher Scientific) in DBB for 3h at RT. Images were acquired using a Zeiss Confocal microscope under identical settings for all experiments conditions.

### RNAscope Multiplex Fluorescent In Situ Hybridization

RNAscope Multiplex Fluorescent Reagent Kits v2 (Advanced Cell Diagnostics, ACD) were used following the manufacturer’s protocols. RNA probes were purchased from ACD Bio-techne for key genes, including CUX2 (Hs-CUX2, 425581; target region 218-1465), CDH7 (Hs-CDH7, 1610931-C1; target region 901-1865), CDH20 (Hs-CDH20, 1610541-C2; target region 901–1865), ADARB2 (Hs-ADARB2, 511651-C3; target region 710– 2219), EPHA7 (Hs-EPHA7, 464661-C2; target region 2–1051) and CADPS2 (Hs-CADPS2, 592711; target region 539 – 1990). Experiments were performed on 16um PFA-fixed tissue sections. Opal fluorophores (520, 570 and 690, Akoya Biosciences) were used for multiplex detection. RNAscope immunohistochemistry was performed without the target retrieval step and primary antibody concentrations were doubled to enhance signal detection.

## RESOURCE AVAILABILITY

### Lead contact

Further information and requests for resources should be directed to and will be fulfilled by the lead contact, Arnold Kriegstein.

### Materials availability

This study did not generate new unique reagents.

## AUTHOR CONTRIBUTIONS

A.K., O.A.B., T.J.N., M.Pa., X.P., E.E.C., A.A.-B., D.R., E.J.H. conceived the project and secured the funding. A.K., O.A.B., and T.J.N., supervised the project. T.M., J.Ag., R.L., C.S., J.C., A.R. prepared samples, performed QC, and generated 10X Multiome libraries. S.W., Q.B. provided assistance with experiments. J.T.H.L., F.M., E.J., Y.C., S.Mk., L.J., M.Pr., L.T., K.T., V.B.P., H.M.C., J.Ha., B.R., O.A.B. generated Xenium spatial transcriptomics datasets. T.M., I.L.L. and G.Z., performed RNAscope and immunostaining experiments. J.M. performed bioinformatics analyses related to 10X Multiome data, with contributions from J.As., Y.H. and J.Ag.. J.T.H.L, F.M. E.J., Y.C., S.Mk., L.J. performed bioinformatics analyses related to Xenium spatial transcriptomics. F.T. provided bioinformatic assistance. J.M., J.H.T.L., J.As. and T.M. prepared figures. J.M. wrote the manuscript with the help of J.H.T.L. and O.A.B. and input from the authors. A.K., O.A.B. and T.J.N. reviewed and edited the manuscript.

## ACKNOWLEDGEMENTS

We thank the members of the Principal Investigators’ research groups involved in this multi-institutional project for their valuable feedback and contributions over the years. We also acknowledge the researchers who participated in the PO-1 network sessions for their insightful discussions and input. This work was supported by the Chan Zuckerberg Initiative. We are grateful to the support staff at all participating institutions, as well as to the team at the Chan Zuckerberg Initiative for their assistance throughout the project. This work was supported by funding from the European Research Council Advanced Grant (789054 to D.H.R.); the Wellcome Trust (to D.H.R. and D.Y.); NIH (P01 NS083513 to A.A.B. D.H.R., X.P., A.R.K.); NIHR Cambridge Biomedical Research Centre (NIHR203312 to D.H.R.); and the Dr. Miriam and Sheldon G. Adelson Medical Research Foundation (to D.H.R.). Views expressed are those of the authors and not necessarily those of the NIHR or the Department of Health and Social Care. T.M. was partly supported by the ANRF Ramanujan fellowship. C.S. acknowledges a PhD fellowship from the Boehringer Ingelheim Fonds. E.J.H acknowledges funding support from P01 NS083513 Neuropathology Core.

## Supplemental Figures

**Figure S1.** (A) Selected examples of fresh frozes samples used in this study. (B) Selected examples of Nissl stainings across Brodmann areas and two time points. (C) Selected quality metrics for snMultiome dataset across 41 samples. (D) UMAP embedding with sample grouping. (E) UMAP embedding with developmental stage (Age) grouping. (F) UMAP embedding with Brodmann area grouping.

**Figure S2.** (A) Hicat clusters are well-represented across individuals and areas at each developmental timepoint. Left: proportions of clusters across individuals. Right: proportions of clusters across Brodmann areas. Dashed vertical lines represent proportions of all cells across individuals. Clusters are sorted from top to bottom by increasing Jensen-Shannon divergence of the cluster distribution from the distribution of all cells across individuals.

**Figure S3.** (A) Same as Figure S2 but for childhood stage after cluster label transfer (see Methods).

**Figure S4.** Same as Figure S2 and Figure S3 but for infancy stage after cluster label transfer (see Methods).

**Figure S5.** (A) Max confidence of label transfer across developmental timepoints for each cluster, in GABAergic neighborhood 1. (B) Max confidence of label transfer across developmental timepoints for each cluster, in GABAergic neighborhood 2. Box plot represents distribution of max confidence across cells.

**Figure S6.** (A) Max confidence of label transfer across developmental timepoints for each cluster, in IT neighborhood. (B) Max confidence of label transfer across developmental timepoints for each cluster, in non-IT neighborhood. (C) Max confidence of label transfer across developmental timepoints for each cluster, in non- neuronal neighborhood. Box plot represents distribution of max confidence across cells.

**Figure S7.** (A) Cell proportion analysis comparing Broca’s area and Motor cortex at each developmental stage at the subclass level. (B) Cell proportion analysis comparing childhood and adult developmental stages in Broca’s area and Motor cortex. (C) Cell proportion analysis comparing Broca’s area and Motor cortex at each developmental stage at taxonomy cluster level.

**Figure S8.** (A) Cell proportion analysis comparing infancy and childhood in Broca’s area (left) and Motor cortex (right) at the cell taxonomy cluster level. (B) Cell proportion analysis comparing childhood and adulthood in Broca’s area (left) and Motor cortex (right) at the cell taxonomy cluster level. No significant results were detected for comparisons in (A) and (B).

**Figure S9.** (A) Distribution of cell area (um sq; left), transcript per cell (middle) and genes per cell (right) per section for Xenium 460-gene panel dataset. (B) Distribution of cell area (um sq; left), transcript per cell (middle) and genes per cell (right) per section for Xenium 5000-gene panel dataset.

**Figure S10.** (A) Selected marker genes for our consensus cell taxonomy in ST datasets. (B) UMAP embeddings for visualization of class, subclass and taxonomy clusters. (C) Banksy-based spatial domain identification across Brodmann areas.

**Figure S11.** (A) Spatial maps of (left to right) cortical layers and area enriched *SST CUX2, VIP CLSTNS and LAMP5 LHX6* neurons in adult BA06 and BA45. (B) Differential cell abundances between BA06 and BA45 areas per developmental stage. Serial Xenium sections per sample were treated as technical replicates. Colors indicate significance after Benjamini-Hochberg adjustment for multiple comparisons (p < 0.05).

**Figure S12.** (A) Spatial maps of (left to right) cortical layers and area enriched all Astrocytes, Astrocyte *GFAP+ GRIA1+,* Astrocyte *GFAP+TNC+,* Astrocyte *HPSE2+GJA1+ and* Oligodendrocyte *ST18+MOBP+* across, infancy, childhood, adolescence and adult BA06 and BA45.

**Figure S13.** (A) Biological functions assigned to time-dependent differentially expressed genes shared between Broca’s area and motor cortex. (B) Selected examples of significant time-dependent DEGs in both areas (with the exception of *SFRP1* that only passes significance threshold in Broca’s area). Significance (*) was considered if adjusted p-value < 0.05. All p-values can be found in Table S5.

**Figure S14.** (A) Spatial maps of *L2-3IT CUX2 LINC00507, PVALB COL15A1, SST FREM1, SST CALB1, SNCG CADPS2,* and *CDH7* expression in BA06 and BA45 at infancy in 5,000-plex Xenium data. Dotplot showing *CDH7* expression in L2-3IT, PVALB, SST and SNCG subclasses in 5,000-plex Xenium data. (B) Spatial maps of *SNCG* subclass in BA06 and BA45 across infancy, childhood, adolescence and adult in 460- plex Xenium data. (C) Spatial maps of *PVALB* subtypes in BA06 and BA45 at adult in 460-plex Xenium data. Dotplot showing CADPS2 expression in *PVALB* subtypes in 460-plex Xenium data.

**Figure S15.** (A–C) Representative RNAscope images showing the expression of *CDH7* (green), *CUX1* (red), *PVALB* (PV) (cyan), DAPI for nuclear staining (blue) in 5-month-old human cortical tissue sections from BA6 (A), BA44 (B), and BA45 (C). For each region, low-magnification tile scans and corresponding higher- magnification images are shown. White and yellow arrowheads indicate CDH7-, CUX1-, and PVALB-positive cells, respectively. UL, upper layer; ML, middle layer; DL, deep layer. Scale bars are indicated in the images.

**Figure S16.** (A–C) Representative RNAscope images showing the expression of *CUX1* (green), *CDH20* (red), and *PVALB* (PV) (cyan) in 5-month-old human cortical tissue sections from BA6 (A), BA44 (B), and BA45 (C). For each region, low-magnification tile scans and corresponding higher-magnification images are shown. White and yellow arrowheads indicate *CUX1*-, *CDH20*-, and *PVALB*-positive cells, respectively. UL, upper layer; ML, middle layer; DL, deep layer. Scale bars are indicated in the images.

**Figure S17.** (A) CellChat analysis capturing *EPHA7*-mediated cell-cell interactions in Broca’s area (left) and motor cortex (right).

**Figure S18.** (A–C) Representative RNAscope images showing the expression of *CADPS2* (green), *PVALB* (PV) (red), and *CUX1* (PV) (cyan) in 4-year-old and 36-year-old human cortical tissue sections from BA6 (A), BA44 (B), and BA44 (C). For each region, low-magnification tile scans and corresponding higher- magnification images are shown. White and yellow arrowheads indicate *CADPS2*-, *PVALB*-, and *CUX1*- positive cells, respectively. UL, upper layer; ML, middle layer; DL, deep layer. Scale bars are indicated in the images.

**Figure S19.** (A) GRNs AUC scores were used to perform dimensionality reduction and obtain a UMAP-based embedding, which reflects separation of Subclasses. (B) Selected GRNs governed by TFs acting as repressors, enriched in different subclasses and summarized by transcription factor expression, region-based AUC scores, and gene-based AUC scores. (C) Number of SCENIC+ GRNs showing differential activity between Broca’s area and motor cortex, per Subclass.

**Figure S20.** (A) *MSX2* expression largely restricted to Astrocyte *GFAP+GRIA1+* and *GFAP+TNC+* in our consensus cell taxonomy. (B) Spatial maps of *GFAP+GRIA1+* astrocytes in BA06 and BA45 at infancy (top), childhood (middle) and adulthood (bottom) in Xenium 5,000-plex data. Histograms show normalized abundance per cortical depth bin (25 bins). (C) Dotplot shows HS3ST3A1 expression in Astrocyte *GFAP+GJA1+*, Astrocyte *GFAP+GRIA1+* and Astrocyte *GFAP+TNC+* between motor (BA06) and Broca’s (BA45) areas in 5,000-plex Xenium data.

**Figure S21.** (A) Standardized disease relevance score across stages per Subclass and cortical area. (B) Area- specific subclass disease relevance score enrichment for ASD, ADHD, SCZ, and ALZ at the cell cluster taxonomy level, per developmental stage and cortical area. *SST PROM1* and *SST CUX2* interneuron subtypes are highlighted as implicated across areas in ASD/ADHD at pediatric stages. Significance for subclass enrichment in disease relevance scores was considered if FDR < 0.05 and odds ratio > 1.4 (two-sided Fisher’s exact tests with Benjamini–Hochberg correction).

**Figure S22.** (A) Per-cell-type sign-concordance between BA22 verbal-vs-non-verbal condition and Broca’s area vs motor cortex differential expression, across matched cell types and developmental stages. Dot size: number of concordant genes; color: −log10 one-sided binomial p for excess concordance (clipped at 4). Concordance is confined to L4/5 IT, L2/3 IT and PVALB neurons at childhood (B) Gene-level comparison of the two contrasts in L4/5 IT (left), L2/3 IT (middle) and PVALB (right) neurons, at childhood. Each point is a gene DE in both contrasts (p < 0.05); x, Broca − motor log2FC; y, verbal − non-verbal log2FC. Red, sign-concordant (Q1/Q3); blue, opposite (Q2/Q4); quadrant counts annotated. One-sided binomial test: L4/5 IT 95/126 (75%, p = 4.8 × 10^-9^), driven by genes up-regulated in both (Q1), L2/3 IT 87/114 (76%, p = 7.6 × 10^-9^), driven by genes down- regulated in both (Q3); PVALB 60/86 (70%, p = 1.6 × 10^-^⁴), driven by genes up-regulated in both (Q1).

## Tables

**Table S1.** Sample-level metadata of 10X Multiome dataset across Brodmann areas.

**Table S2.** Subclass-level summary of 10X Multiome dataset across Brodmann areas.

**Table S3.** Taxonomy-level summary of 10X Multiome dataset across Brodmann areas.

**Table S4.** Cell compositional analysis results, summarized from both subclass- and cluster-level analyses.

**Table S5.** Pseudobulk-based linear mixed model analysis results.

**Table S6.** Gene ontology and semantic similarity analysis results for time-dependent DE genes.

**Table S7.** SCENIC+ Gene regulatory network metrics.

**Table S8.** Sign concordance test on DEGs from BA22 and Broca’s area vs Motor cortex.

**Table S9.** Significant differentially accessible regions identified, with Subclass and developmental stage information.

**Table S10.** Significant differentially accessible regions that overlap with evolutionary-relevant genomic regions, with GREAT-based gene assignment.

## REFERENCES

1. Darwin, C., Desmond, A. J. & Moore, J. R. The Descent of Man, and Selection in Relation to Sex. (Penguin, London, 2004).

2. Ziegenfusz, S., Paynter, J., Flückiger, B. & Westerveld, M. F. A systematic review of the academic achievement of primary and secondary school-aged students with developmental language disorder. Autism Dev. Lang. Impair. 7, 23969415221099396 (2022).

3. Yew, S. G. K. & O’Kearney, R. Emotional and behavioural outcomes later in childhood and adolescence for children with specific language impairments: Meta-analyses of controlled prospective studies. J. Child Psychol. Psychiatry 54, 516–524 (2013).

4. Poeppel, D. The neuroanatomic and neurophysiological infrastructure for speech and language. Curr. Opin. Neurobiol. 28, 142–149 (2014).

5. Wolna, A. et al. The extended language network: Language-responsive brain areas whose contributions to language remain to be discovered. bioRxiv 2025.04.02.646835 (2026) doi:10.1101/2025.04.02.646835.

6. Broca, P. Remarques sur le siège de la faculté du langage articulé, suivies d’une observation d’aphémie (perte de la parole). Bulletin et mémoires de la Société Anatomique de Paris 6, 330–357 (1861).

7. Eggert Gertrude H. & Eggert Gertrude H. Wernicke’s Works on Aphasia : A Sourcebook and Review. Volume 1, Early Sources in Aphasia and Related Disorders (De Gruyter Mouton, Berlin;, 2019).

8. Lichteim, L. On Aphasia. Brain 7, 433–484 (1885).

9. Geschwind, N. The Organization of Language and the Brain. Science (1979). 170, 940–944 (1970).

10. Brodmann, K. & Gary, L. J. Brodmann’s Localization in the Cerebral Cortex : The Principles of Comparative Localisation in the Cerebral Cortex Based on (Springer, New York, NY, 2006).

11. Amunts, K. & Zilles, K. Architecture and organizational principles of Broca’s region. Trends Cogn. Sci. 16, 418–426 (2012).

12. Zilles, K. & Amunts, K. Cytoarchitectonic and receptorarchitectonic organization in Broca’s region and surrounding cortex. Curr. Opin. Behav. Sci. 21, 93–105 (2018).

13. Palomero-Gallagher, N. & Zilles, K. Differences in cytoarchitecture of Broca’s region between human, ape and macaque brains. Cortex 118, 132–153 (2019).

14. Wang, Z. et al. Transcriptomics and functional genomics implicate WNT3 in hemispheric lateralization of speech production. iScience 29, 114692 (2026).

15. Khaitovich, P. et al. Regional Patterns of Gene Expression in Human and Chimpanzee Brains. Genome Res. 14, 1462 (2004).

16. Wong, M. M. K. et al. The neocortical infrastructure for language involves region-specific patterns of laminar gene expression. Proc. Natl. Acad. Sci. U. S. A. 121, e2401687121 (2024).

17. Wagstyl, K. et al. Transcriptional Cartography Integrates Multiscale Biology of the Human Cortex. Elife 12, (2024).

18. Zilles, K., Bacha-Trams, M., Palomero-Gallagher, N., Amunts, K. & Friederici, A. D. Common molecular basis of the sentence comprehension network revealed by neurotransmitter receptor fingerprints. Cortex 63, 79–89 (2015).

19. Palomero-Gallagher, N. & Zilles, K. Cortical layers: Cyto-, myelo-, receptor- and synaptic architecture in human cortical areas. Neuroimage 197, 716–741 (2019).

20. Bakken, T. E. et al. Comparative cellular analysis of motor cortex in human, marmoset and mouse. Nature 2021 598:7879 598, 111–119 (2021).

21. Petukhov, V. et al. Case-control analysis of single-cell RNA-seq studies. bioRxiv 2022.03.15.484475 (2022) doi:10.1101/2022.03.15.484475.

22. Brauer, J., Anwander, A., Perani, D. & Friederici, A. D. Dorsal and ventral pathways in language development. Brain Lang. 127, 289–295 (2013).

23. Dubois, J. et al. Exploring the Early Organization and Maturation of Linguistic Pathways in the Human Infant Brain. Cerebral Cortex 26, 2283–2298 (2016).

24. Tucić, M., Stamenković, V. & Andjus, P. The Extracellular Matrix Glycoprotein Tenascin C and Adult Neurogenesis. Front. Cell Dev. Biol. 9, 674199 (2021).

25. Cooper, M. L. et al. Astrocytes connect specific brain regions through plastic networks. Nature 2026 655:8121 655, 183–191 (2026).

26. Shiu, F. H. et al. ADGRB1 contributes to astrocyte-mediated phagocytosis of excitatory synapses. Exp. Neurol. 395, 115451 (2026).

27. Hoffman, G. E., et al. Efficient differential expression analysis of large-scale single cell transcriptomics data using dreamlet. bioRxiv 2023.03.17.533005 (2024) doi:10.1101/2023.03.17.533005.

28. Moissidis, M. et al. A postnatal molecular switch drives activity-dependent maturation of parvalbumin interneurons. Cell 188, 5555–5575.e26 (2025).

29. Redies, C. Cadherins in the central nervous system. Prog. Neurobiol. 61, 611–648 (2000).

30. Duan, X., Krishnaswamy, A., De La Huerta, I. & Sanes, J. R. Type II cadherins guide assembly of a direction-selective retinal circuit. Cell 158, 793–807 (2014).

31. Williams, M. E. et al. Cadherin-9 Regulates Synapse-Specific Differentiation in the Developing Hippocampus. Neuron 71, 640–655 (2011).

32. Jézéquel, J. et al. Cadherins orchestrate specific patterns of perisomatic inhibition onto distinct pyramidal cell populations. Nature Communications 2025 16:1 16, 4481- (2025).

33. Jin, S., Plikus, M. V. & Nie, Q. CellChat for systematic analysis of cell–cell communication from single-cell transcriptomics. Nature Protocols 2024 20:1 20, 180–219 (2024).

34. Lévy, J. et al. EPHA7 haploinsufficiency is associated with a neurodevelopmental disorder. Clin. Genet. 100, 396–404 (2021).

35. Luo, X. et al. 3D Genome of macaque fetal brain reveals evolutionary innovations during primate corticogenesis. Cell 184, 723–740.e21 (2021).

36. Okbay, A. et al. Polygenic prediction of educational attainment within and between families from genome-wide association analyses in 3 million individuals. Nature Genetics 2022 54:4 54, 437–449 (2022).

37. Mulatinho, M. V. et al. Severe intellectual disability, omphalocele, hypospadia and high blood pressure associated to a deletion at 2q22.1q22.3: case report. Molecular Cytogenetics 2012 5:1 5, 30- (2012).

38. Smigiel, R. et al. Severe clinical course of Hirschsprung disease in a Mowat-Wilson syndrome patient. Journal of Applied Genetics 2010 51:1 51, 111–113 (2010).

39. Kutsche, K. et al. Mutations in ARHGEF6, encoding a guanine nucleotide exchange factor for Rho GTPases, in patients with X-linked mental retardation. Nature Genetics 2000 26:2 26, 247–250 (2000).

40. Davies, G. et al. Genome-wide association studies establish that human intelligence is highly heritable and polygenic. Molecular Psychiatry 2011 16:10 16, 996–1005 (2011).

41. Benyamin, B. et al. Childhood intelligence is heritable, highly polygenic and associated with FNBP1L. Molecular Psychiatry 2014 19:2 19, 253–258 (2013).

42. Ziegler, D. V. et al. Calcium channel ITPR2 and mitochondria–ER contacts promote cellular senescence and aging. Nature Communications 2021 12:1 12, 720- (2021).

43. van Es, M. A. et al. ITPR2 as a susceptibility gene in sporadic amyotrophic lateral sclerosis: a genome- wide association study. Lancet Neurol. 6, 869–877 (2007).

44. Vernot, B. et al. Excavating Neandertal and Denisovan DNA from the genomes of Melanesian individuals. Science (1979). 352, 235–239 (2016).

45. Peyrégne, S. et al. Detecting ancient positive selection in humans using extended lineage sorting. Genome Res. 27, 1563–1572 (2017).

46. Bravo González-Blas, C., et al. SCENIC+: single-cell multiomic inference of enhancers and gene regulatory networks. Nature Methods 2023 20:9 20, 1355–1367 (2023).

47. Aibar, S. et al. SCENIC: single-cell regulatory network inference and clustering. Nature Methods 2017 14:11 14, 1083–1086 (2017).

48. Harris, H. K. et al. Disruption of RFX family transcription factors causes autism, attention- deficit/hyperactivity disorder, intellectual disability, and dysregulated behavior. Genetics in Medicine 23, 1028–1040 (2021).

49. Fu, J. M. et al. Rare coding variation provides insight into the genetic architecture and phenotypic context of autism. Nature Genetics 2022 54:9 54, 1320–1331 (2022).

50. Suresh, V. et al. Molecular dynamics of Brodmann Area 22 in development and autism. bioRxiv 2026.03.31.715694 (2026) doi:10.64898/2026.03.31.715694.

51. Stromme, P., Mangelsdorf, M. E., Scheffer, I. E. & Gécz, J. Infantile spasms, dystonia, and other X-linked phenotypes caused by mutations in Aristaless related homeobox gene, ARX. Brain Dev. 24, 266–268 (2002).

52. Kato, M., Das, S., Petras, K., Sawaishi, Y. & Dobyns, W. B. Polyalanine expansion of ARX associated with cryptogenic West syndrome. Neurology 61, 267–268 (2003).

53. Akula, S. K. et al. The spectrum of movement disorders in young children with ARX -related epilepsy- dyskinesia syndrome. Ann. Clin. Transl. Neurol. 11, 1643 (2024).

54. Strømme, P. et al. Mutations in the human ortholog of Aristaless cause X-linked mental retardation and epilepsy. Nature Genetics 2002 30:4 30, 441–445 (2002).

55. Bienvenu, T. et al. ARX, a novel Prd-class-homeobox gene highly expressed in the telencephalon, is mutated in X-linked mental retardation. Hum. Mol. Genet. 11, 981–991 (2002).

56. Meng, J. et al. Single-cell spatial map of cis-regulatory elements for disease-related genes in the macaque cortex. Nature Communications 2026 17:1 17, 4041- (2026).

57. Chen, A. et al. Single-cell spatial transcriptome reveals cell-type organization in the macaque cortex. Cell 186, 3726–3743.e24 (2023).

58. Tecle, E., Diaz-Balzac, C. A. & Bülow, H. E. Distinct 3-O-Sulfated Heparan Sulfate Modification Patterns Are Required for kal-1−Dependent Neurite Branching in a Context-Dependent Manner in Caenorhabditis elegans. G3 Genes|Genomes|Genetics 3, 541–552 (2013).

59. Maïza, A. et al. 3-O-sulfated heparan sulfate interactors target synaptic adhesion molecules from neonatal mouse brain and inhibit neural activity and synaptogenesis in vitro. Scientific Reports 2020 10:1 10, 19114- (2020).

60. Jacot-Descombes, S. et al. Decreased pyramidal neuron size in Brodmann areas 44 and 45 in patients with autism. Acta Neuropathologica 2012 124:1 124, 67–79 (2012).

61. Ibrahim, K. et al. Reduced Amygdala–Prefrontal Functional Connectivity in Children With Autism Spectrum Disorder and Co-occurring Disruptive Behavior. Biol. Psychiatry Cogn. Neurosci. Neuroimaging 4, 1031–1041 (2019).

62. Bak, T. H., O’Donovan, D. G., Xuereb, J. H., Boniface, S. & Hodges, J. R. Selective impairment of verb processing associated with pathological changes in Brodmann areas 44 and 45 in the motor neurone disease–dementia–aphasia syndrome. Brain 124, 103–120 (2001).

63. Yu, F. et al. Variant to function mapping at single-cell resolution through network propagation. Nature Biotechnology 2022 40:11 40, 1644–1653 (2022).

64. Duncan, L. E. et al. Mapping the cellular etiology of schizophrenia and complex brain phenotypes. Nature Neuroscience 2025 28:2 28, 248–258 (2025).

65. De La Torre-Ubieta, L., Won, H., Stein, J. L. & Geschwind, D. H. Advancing the understanding of autism disease mechanisms through genetics. Nature Medicine 2016 22:4 22, 345–361 (2016).

66. Nott, A. et al. Brain cell type–specific enhancer–promoter interactome maps and disease-risk association. Science (1979). 366, 1134–1139 (2019).

67. Rohden, F. et al. Glial reactivity correlates with synaptic dysfunction across aging and Alzheimer’s disease. Nature Communications 2025 16:1 16, 5653- (2025).

68. Wamsley, B. et al. Molecular cascades and cell type–specific signatures in ASD revealed by single-cell genomics. Science (1979). 384, (2024).

69. Velmeshev, D. et al. Single-cell genomics identifies cell type–specific molecular changes in autism. Science (1979). 364, 685–689 (2019).

70. Geschwind, D. H. Genetics of autism spectrum disorders. Trends Cogn. Sci. 15, 409–416 (2011).

71. Xue, J. R. et al. The functional and evolutionary impacts of human-specific deletions in conserved elements. Science (1979). 380, (2023).

72. Whalen, S. et al. Three-dimensional genome rewiring in loci with human accelerated regions. Science (1979). 380, (2023).

73. Mangan, R. J. et al. Adaptive sequence divergence forged new neurodevelopmental enhancers in humans. Cell 185, 4587–4603.e23 (2022).

74. Peyregne, S., Boyle, M. J., Dannemann, M. & Prufer, K. Detecting ancient positive selection in humans using extended lineage sorting. Genome Res. 27, 1563–1572 (2017).

75. Casten, L. G. et al. Ancient regulatory evolution shapes individual language abilities in present-day humans. Sci. Adv. 12, (2026).

76. McLean, C. Y. et al. GREAT improves functional interpretation of cis-regulatory regions. Nat. Biotechnol. 28, 495–501 (2010).

77. Tanigawa, Y., Dyer, E. S. & Bejerano, G. WhichTF is functionally important in your open chromatin data? PLoS Comput. Biol. 18, (2022).

78. Gehman, L. T. et al. The splicing regulator Rbfox2 is required for both cerebellar development and mature motor function. Genes Dev. 26, 445 (2012).

79. Jacko, M. et al. Rbfox Splicing Factors Promote Neuronal Maturation and Axon Initial Segment Assembly. Neuron 97, 853–868.e6 (2018).

80. Gialluisi, A. et al. Genome-wide screening for DNA variants associated with reading and language traits. Genes Brain Behav. 13, 686–701 (2014).

81. Gialluisi, A., Guadalupe, T., Francks, C. & Fisher, S. E. Neuroimaging genetic analyses of novel candidate genes associated with reading and language. Brain Lang. 172, 9–15 (2017).

82. Smith, R. S. et al. Early role for a Na+,K+-ATPase (ATP1A3) in brain development. Proc. Natl. Acad. Sci. U. S. A. 118, e2023333118 (2021).

83. Smedemark-Margulies, N., et al. A novel de novo mutation in ATP1A3 and childhood-onset schizophrenia. Molecular Case Studies 2, a001008 (2016).

84. Heinzen, E. L. et al. De novo mutations in ATP1A3 cause alternating hemiplegia of childhood. Nature Genetics 2012 44:9 44, 1030–1034 (2012).

85. Lenneberg, E. H. The Biological Foundations of Language. Hosp. Pract. 2, 59–67 (1967).

86. Werker, J. F. & Hensch, T. K. Critical periods in speech perception: New directions. Annu. Rev. Psychol. 66, 173–196 (2015).

87. Fisher, S. E. & Vernes, S. C. Genetics and the Language Sciences. Annu. Rev. Linguist. 1, 289–310 (2015).

88. Tsyporin, J. et al. Competing programs shape cortical sensorimotor–association axis development. Nature 2026 1–12 (2026) doi:10.1038/s41586-026-10699-x.

89. Matsunaga, E. et al. Dynamic Expression of Cadherins Regulates Vocal Development in a Songbird. PLoS One 6, e25272 (2011).

90. Matsunaga, E. & Okanoya, K. Expression analysis of cadherins in the songbird brain: Relationship to vocal system development. Journal of Comparative Neurology 508, 329–342 (2008).

91. Matsunaga, E. & Okanoya, K. Cadherins: potential regulators in the faculty of language. Curr. Opin. Neurobiol. 28, 28–33 (2014).

92. Green, R. E. et al. A draft sequence of the neandertal genome. Science (1979). 328, 710–722 (2010).

93. Aprile, D. et al. Benchmarking cerebellar organoids to model autism spectrum disorder and human brain evolution. bioRxiv 2025.05.14.653684 (2025) doi:10.1101/2025.05.14.653684.

94. Choe, H. N. et al. Estrogen and sex-dependent loss of the vocal learning system in female zebra finches. Horm. Behav. 129, 104911 (2021).

95. Pfenning, A. R. et al. Convergent transcriptional specializations in the brains of humans and song- learning birds. Science (1979). 346, (2014).

96. Wirthlin, M. E. et al. Vocal learning-associated convergent evolution in mammalian proteins and regulatory elements. Science (1979). 383, (2024).

97. Arnon, I. et al. What enables human language? A biocultural framework. Science (1979). 390, (2025).

98. Clarke, L. E. & Barres, B. A. Emerging roles of astrocytes in neural circuit development. Nature Reviews Neuroscience 2013 14:5 14, 311–321 (2013).

99. Shimizu, T. et al. Oligodendrocyte dynamics dictate cognitive performance outcomes of working memory training in mice. Nat. Commun. 14, (2023).

100. Wong, F. K. & Marín, O. Developmental Cell Death in the Cerebral Cortex. Annu. Rev. Cell Dev. Biol. 35, 523–542 (2019).

101. Ponce de León, M. S., et al. The primitive brain of early Homo. Science (1979). 372, (2021).

102. Young, M. D. & Behjati, S. SoupX removes ambient RNA contamination from droplet-based single-cell RNA sequencing data. Gigascience 9, (2020).

103. Germain, P. L., Robinson, M. D., Lun, A., Garcia Meixide, C. & Macnair, W. Doublet identification in single-cell sequencing data using *scDblFinder*. F1000Research 2022 10:979 10, 979 (2022).

104. Thibodeau, A. et al. AMULET: a novel read count-based method for effective multiplet detection from single nucleus ATAC-seq data. Genome Biol. 22, 1–19 (2021).

105. Ma, S. et al. Molecular and cellular evolution of the primate dorsolateral prefrontal cortex. Science (1979). 377, (2022).

106. Gabitto, M. I. et al. Integrated multimodal cell atlas of Alzheimer’s disease. Nat. Neurosci. 27, 2366– 2383 (2024).

107. Tasic, B. et al. Shared and distinct transcriptomic cell types across neocortical areas. Nature 2018 563:7729 563, 72–78 (2018).

108. Gao, Y. et al. Continuous cell-type diversification in mouse visual cortex development. Nature 2025 647:8088 647, 127–142 (2025).

109. Szklarczyk, D. et al. The STRING database in 2017: quality-controlled protein–protein association networks, made broadly accessible. Nucleic Acids Res. 45, D362–D368 (2017).

110. Persad, S. et al. SEACells infers transcriptional and epigenomic cellular states from single-cell genomics data. Nature Biotechnology 2023 41:12 41, 1746–1757 (2023).

111. Suo, S. et al. Revealing the Critical Regulators of Cell Identity in the Mouse Cell Atlas. Cell Rep. 25, 1436–1445.e3 (2018).

112. Brionne, A., Juanchich, A. & Hennequet-Antier, C. ViSEAGO: a Bioconductor package for clustering biological functions using Gene Ontology and semantic similarity. BioData Mining 2019 12:1 12, 16- (2019).

113. Bioconductor - topGO. https://bioconductor.org/packages/release/bioc/html/topGO.html.

114. Stuart, T. et al. Comprehensive Integration of Single-Cell Data. Cell 177, 1888–1902.e21 (2019).

115. Rahimi, A., Vale-Silva, L. A., Fälth Savitski, M., Tanevski, J. & Saez-Rodriguez, J. DOT: a flexible multi- objective optimization framework for transferring features across single-cell and spatial omics. Nature Communications 2024 15:1 15, 4994- (2024).

116. Korsunsky, I. et al. Fast, sensitive and accurate integration of single-cell data with Harmony. Nature Methods 2019 16:12 16, 1289–1296 (2019).

117. Singhal, V. et al. BANKSY unifies cell typing and tissue domain segmentation for scalable spatial omics data analysis. Nature Genetics 2024 56:3 56, 431–441 (2024).

118. Büttner, M., Ostner, J., Müller, C. L., Theis, F. J. & Schubert, B. scCODA is a Bayesian model for compositional single-cell data analysis. Nature Communications 2021 12:1 12, 6876- (2021).

119. Yu, F. et al. Variant to function mapping at single-cell resolution through network propagation. Nat. Biotechnol. 40, 1644–1653 (2022).

120. Yang, J., Lee, S. H., Goddard, M. E. & Visscher, P. M. GCTA: a tool for genome-wide complex trait analysis. Am. J. Hum. Genet. 88, 76–82 (2011).

121. Yang, J. et al. Conditional and joint multiple-SNP analysis of GWAS summary statistics identifies additional variants influencing complex traits. Nat. Genet. 44, 369–375 (2012).

122. Squair, J. W. et al. Confronting false discoveries in single-cell differential expression. Nature Communications 2021 12:1 12, 5692- (2021).

123. Gel, B. et al. regioneR: an R/Bioconductor package for the association analysis of genomic regions based on permutation tests. Bioinformatics 32, 289–291 (2016).

